# Cell cycle oscillators underlying orderly proteolysis of E2F8

**DOI:** 10.1101/672964

**Authors:** Danit Wasserman, Sapir Nachum, Meital Cohen, Taylor P Enrico, Meirav Noach-Hirsh, Jasmin Parasol, Sarit Zomer-Polak, Naomi Auerbach, Evelin Sheinberger-Chorni, Hadas Nevenzal, Nofar Levi-Dadon, Xianxi Wang, Roxane Lahmi, Efrat Michaely, Doron Gerber, Michael J. Emanuele, Amit Tzur

**Affiliations:** Faculty of Life Sciences and Institute of Nanotechnology and Advanced Materials, Bar-llan University, Ramat-Gan 5290002, Israel; Lineberger Comprehensive Cancer Center, Department of Pharmacology, The University of North Carolina at Chapel Hill, Chapel Hill, NC 27599, United States of America

## Abstract

E2F8 is a transcriptional repressor that antagonizes the canonical cell cycle transcription factor E2F1. Despite the importance of this atypical E2F family member in cell cycle, apoptosis and cancer, we lack a complete description of the mechanisms that control its dynamics. To address this question, we developed a complementary set of static and dynamic cell-free systems of human origin, which recapitulate inter-mitotic and G1 phases, and a full transition from pro-metaphase to G1. This revealed an interlocking molecular switch controlling E2F8 degradation at mitotic exit, involving dephosphorylation of Cdk1 sites in E2F8 and the activation of APC/C^Cdh1^, but not APC/C^Cdc20^. Further, we revealed a differential stability of E2F8, accounting for its accumulation in late G1 while APC/C^Cdh1^ is still active and suggesting a key role for APC/C in controlling G1-S transcription. Finally, we identified SCF-Cyclin F as the ubiquitin ligase controlling E2F8 in G2-phase. Altogether, our data provide new insights into the regulation of E2F8 throughout the cell cycle, illuminating an extensive coordination between phosphorylation, ubiquitination and transcription in promoting orderly cell cycle progression.

## Introduction

The E2F family of transcription factors plays a pivotal role in regulating pro- and anti-proliferative processes, with implications in tissue homeostasis and human disease, most notably cancer (Chen, Tsai et al., 2009). E2F1, the canonical member of the E2F family, is at the crossroads of the cell cycle and cell death, triggering the gene program dictating S-phase and mitotic entry on the one hand, and, on the other hand, apoptosis (Hallstrom & Nevins, 2009, Polager & Ginsberg, 2008, Thurlings & de Bruin, 2016).

The activity of E2F1 is balanced by two of its direct targets E2F7 and E2F8, also known as atypical E2Fs (Christensen, Cloos et al., 2005, de Bruin, Maiti et al., 2003, Di Stefano, Jensen et al., 2003, Maiti, Li et al., 2005). E2F7 and E2F8 are structurally and functionally related (Lammens, Li et al., 2009); they cooperate in repressing the transcription of E2F1 and its downstream target genes by binding consensus E2F motifs along their promoters. The result is a negative feedback circuit whose dynamics control cell fate, tissue homeostasis and development by mechanisms that are still not entirely clear (Li, Ran et al., 2008, Ouseph, Li et al., 2012). Despite being part of the ‘repressive’ branch of E2F proteins, E2F7 and E2F8 belong to the pro-proliferative gene network underlying cell proliferation (Cohen, Vecsler et al., 2013).

E2F1, as well as E2F7 and E2F8, are regulated post-translationally via temporal proteolysis. The anaphase-promoting complex/cyclosome (APC/C) is a multi-subunit cell cycle ubiquitin ligase and core component of the cell cycle machinery (King, Peters et al., 1995, Sudakin, Ganoth et al., 1995). The APC/C uses two related co-activators termed Cdc20 and Cdh1, which bind substrates and recruit them to the APC/C for ubiquitination and subsequent degradation (Kernan, Bonacci et al., 2018). We previously identified both E2F7 and E2F8 as targets of the Cdh1-bound form of APC/C (APC/C^Cdh1^) (Cohen et al., 2013). These findings, together with other supporting studies (Boekhout, Yuan et al., 2016), shifted the model by which the E2F1-E2F7-E2F8 circuitry communicates with the cell cycle clock to regulate the transition from the G1-phase of the cell cycle into S-phase. Nevertheless, the exact inter-dynamics of E2F1-E2F7-E2F8 circuitry throughout G1 and the mechanism by which they are achieved are not entirely resolved. No less obscure is the interplay between E2F1 and atypical E2Fs during G2-phase and mitosis. Dissecting these complex signaling circuits is important for understanding the decision making mechanisms at two critical points in the life of a proliferating cell – commitment to DNA replication and division.

Cell-free systems are known for their capacity to reproduce complex cellular processes *in vitro* while maintaining a physiologically relevant context, bridging the gap between *in vivo* and *in vitro*. These systems are optimal for direct and quantitative analysis of time-specific molecular events, including phosphorylation and ubiquitination, circumventing caveats associated with long-term *in vivo* manipulations. Cell-free systems can be either ‘static’, meaning they capture a certain physiological state (*e.g*., interphase), or ‘dynamic’, reproducing transitions between phases, including the spatiotemporal dynamics of proteins, chromatin and complex cellular structures (*e.g*., mitotic spindle). Mitotic entry and exit, metaphase-to-anaphase transition and cytokinesis were all demonstrated in frog egg extracts (Funabiki & Murray, 2000, Murray, Desai et al., 1996, Murray & Kirschner, 1989, Nguyen, Groen et al., 2014), as well as in other early embryonic systems (Telley, Gaspar et al., 2012). Mammalian cell-free systems have been gradually integrated in cell cycle research in the last 20 years. Extracts from synchronous cell populations provide a biochemically amenable system that contains all of the critical enzymes necessary to recapitulate and analyze protein degradation, phosphorylation and other key cell cycle signaling events in a somatic 4-stage cell cycle context lacking in egg extracts (Ayad, Rankin et al., 2005, Nguyen, Gitig et al., 1999, Rape & Kirschner, 2004, Rape, Reddy et al., 2006).

Here, we developed and utilized a panel of ‘static’ and ‘dynamic’ human cell-free systems with which we disentangle the mechanism underlying temporal dynamics of the E2F8 protein from pro-metaphase to late S-phase. *In vivo* studies addressing the regulation of E2F8 during G2-phase complete the missing piece of the puzzle.

## Results

### Temporal dynamics of E2F8 *vs.* E2F1 across the cell cycle

An overview of E2F8 and E2F1 protein-levels throughout the cell cycle highlights two important points in the context of this study. First, in synchronous HeLa S3 cell populations released from a nocodazole-induced pro-metaphase block, E2F1 accumulates during early-mid G1 coinciding with the peak of APC/C^Cdh1^ activity (**Fig. 1A**). In fact, E2F1 dynamics appear inversely to that of APC/C^Cdh1^ target Kifc1 when analyzed in 30 to 60 min time resolution (**Fig. 1A and Fig. S1**). The rise of E2F8 commences approximately 2.5-3 h after E2F1 initially appears. Still, E2F8 accumulates prior to the G1-S transition and well before the rise of canonical APC/C targets. As early as 7 hours after release from nocodazole, increasing E2F8 levels are evident, yet cells have not yet entered S-phase based on DNA quantification (**Fig. 1A**). We and others previously showed that APC/C^Cdh1^ prevents E2F8 from accumulating in lock-step with E2F1 during mid G1 (Boekhout et al., 2016, Cohen et al., 2013). Thus, it is unclear how E2F8 levels increase in late G1 while APC/C^Cdh1^ is still active. Secondly, E2F8 levels decrease prior to mitosis through an entirely unknown mechanism (**Fig. 1A and B**). Even under a tight positive regulation by E2F1, it is unlikely that this reduction of E2F8 is controlled only at the transcriptional level. Interestingly, while E2F1 is undetectable in pro-metaphase, E2F8 levels are low but still higher than in early-mid G1-phase (**Fig. 1A**).

**Figure 1:**
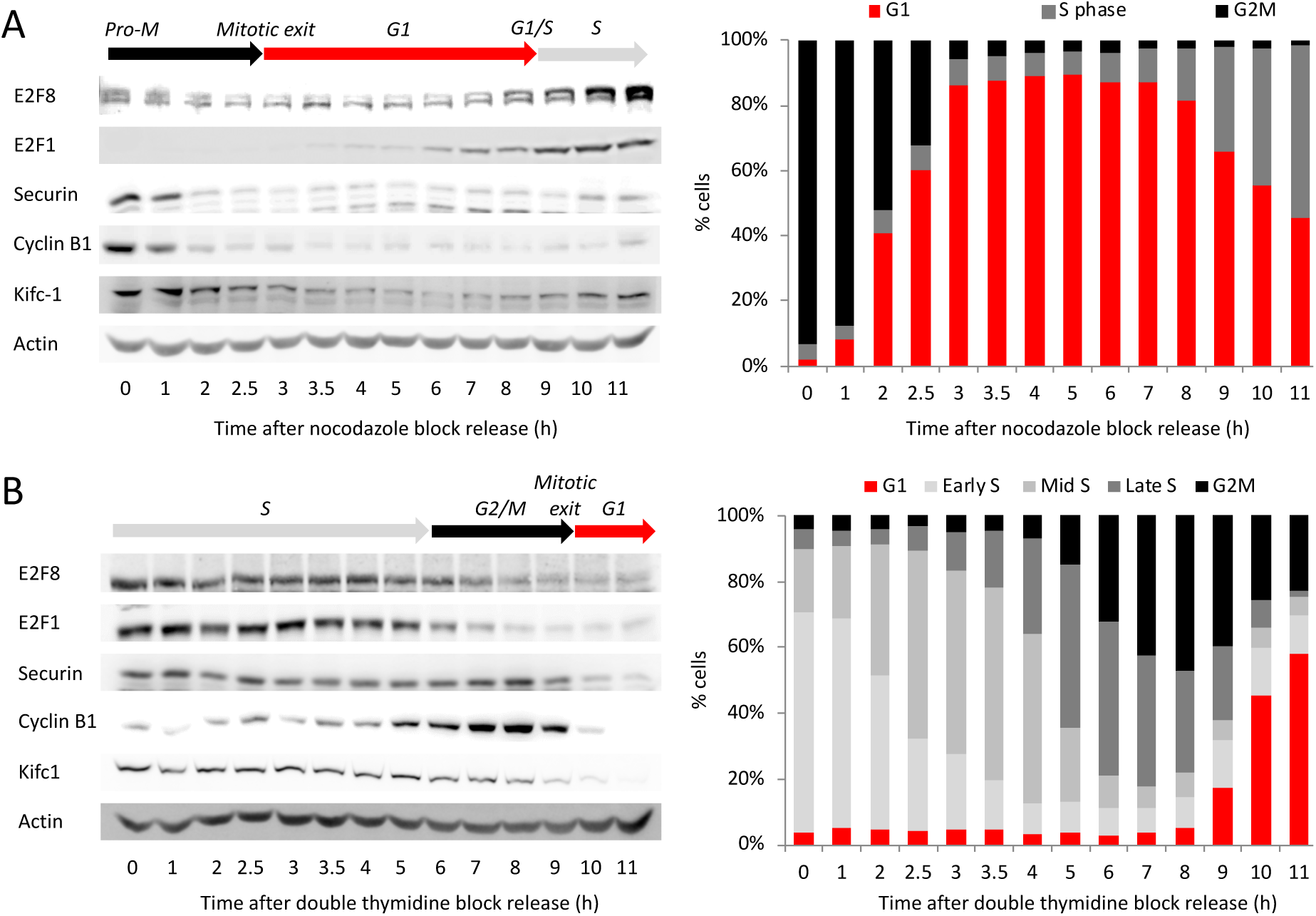
*Temporal dynamics of E2F8* across the cell cycle. Western blot analyses of E2F8, E2F1 and canonical APC/C targets in synchronous S3 cells (DNA distributions are shown). Synchronization methods: release from thymidine-nocodazole block (A) and release from double thymidine block (B).

### Temporal proteolysis of E2F8 in ‘dynamic’ mitotic extracts progressing from pro-metaphase to G1-phase

The activation of APC/C^Cdc20^ following removal of the <u>m</u>itotic <u>c</u>heckpoint <u>c</u>omplex (MCC) and then of APC/C^Cdh1^ (illustrated in **Figure 2A**) are major milestones along mitotic progression and exit that can be monitored by the sequential onset of Securin and Tome-1 degradation at the metaphase-to-anaphase transition and G1 entry, respectively (Ayad, Rankin et al., 2003, Hagting, Den Elzen et al., 2002, Zou, McGarry et al., 1999, Zur & Brandeis, 2001). Mitotic exit is also characterized by a massive dephosphorylation wave orchestrated by the inactivation of Cdk1 and activation of phosphatases during mitotic exit (Grallert, Boke et al., 2015, Powers & Hall, 2017, Wurzenberger & Gerlich, 2011).

**Figure 2:**
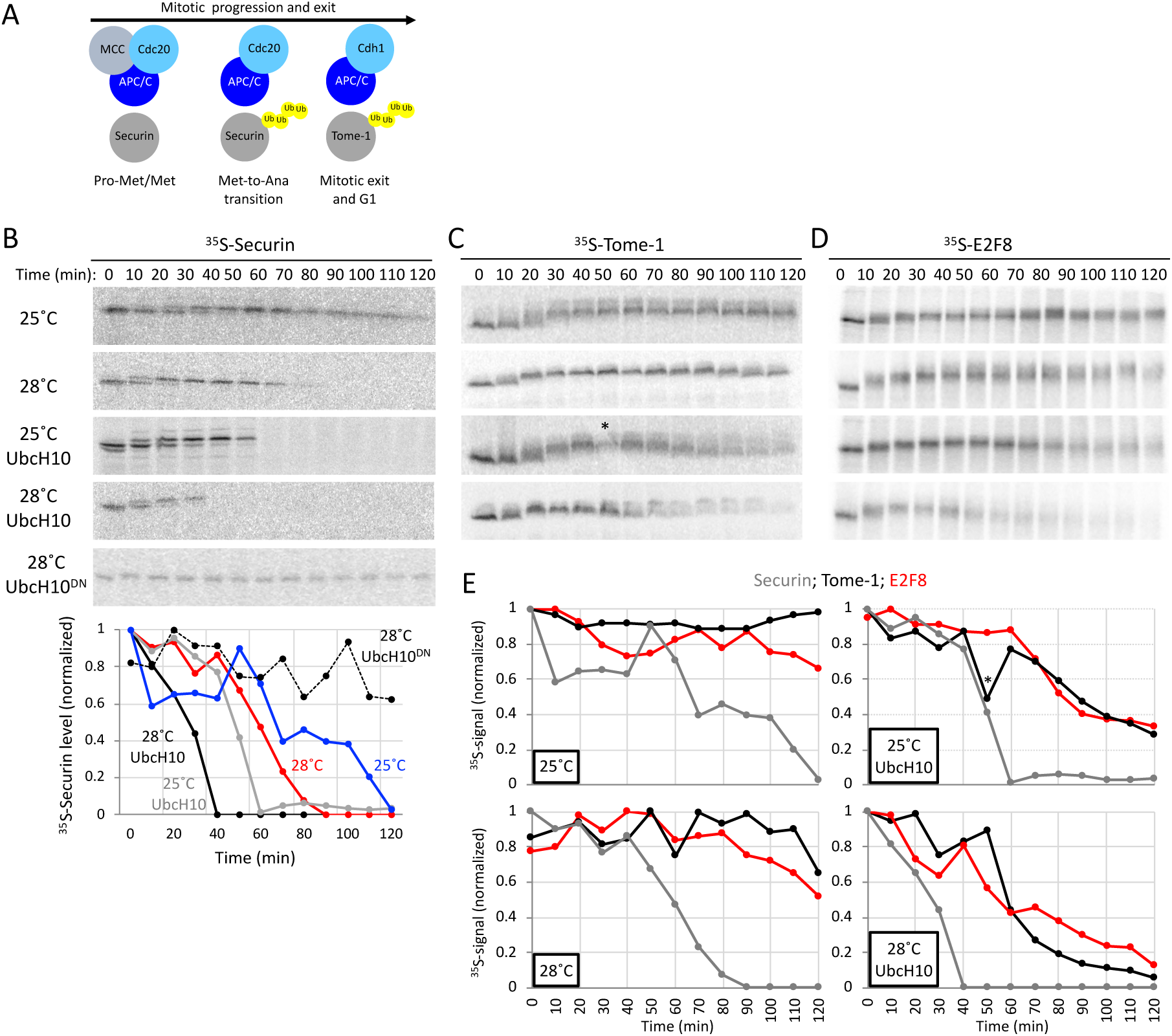
Temporal proteolysis of E2F8 in ‘dynamic’ mitotic extracts progressing from pro-metaphase to G1. **(A)** Schematic of molecular milestones along mitosis. At <u>pro-met</u>aphase (Pro-Met) and <u>met</u>aphase (Met), the <u>m</u>itotic <u>c</u>heckpoint <u>c</u>omplex (MCC) prevents from APC/C^Cdc20^ to ubiquitinate Securin. MCC removal induces Securin and Cyclin B1 degradation and the metaphase-to-anaphase (Met-to-Ana) transition. The drop in Cyclin B1 levels induces i) the switch from APC/C^Cdc20^ to APC/C^Cdh1^ activity; ii) the degradation of Tome-1 and other Cdh1-sepcific targets; and iii) mitotic exit into G1. **(B - E)** Cell extracts were made from thymidine/nocodazole-arrested S3 cells. MCC removal and mitotic exit occur spontaneously in these extracts in a temperature-controlled manner or by adding recombinant UbcH10. The addition of dominant negative UbcH10 (UbcH10^DN^) blocks mitotic progression and exit (B). Temporal proteolysis and (de)phosphorylation-induced electrophoretic mobility shifts of Securin and Tome-1 are shown (B and C). At optimal reaction condition (UbcH10; 28°C), extracts eventually reach a G1-like state in which APC/C^Cdh1^ is active (C). Time-dependent degradations of Securin (B), Tome-1 (C) and E2F8 (D) [^35^S-labeled *<u>i</u>n <u>v</u>itro* translated (IVT) products] were assayed by SDS-PAGE and autoradiography. Source data (B-D) and quantification (E) are shown (* marks a deformed band). Temporal proteolysis and electrophoretic mobility shifts of E2F8 and Tome-1 are highly similar.

Pro-metaphase extracts can be made from nocodazole-arrested HeLa S3 (S3) cells (Ayad et al., 2005, Rape & Kirschner, 2004). Here we show that this system could be dynamically controlled to recreate the biochemistry and signaling of progression from pro-metaphase to G1-phase. Spindle checkpoint removal occurs spontaneously in pro-metaphase extracts in a temperature-controlled manner or, more efficiently, by adding the APC/C E2 enzyme UbcH10. On the flip side, dominant negative UbcH10 (UbcH10^DN^) blocks the extracts, making it ‘static’ in a pro-metaphase-like state for hours. These features are manifested by the temporal proteolysis and Cdk1-induced electrophoretic mobility shift of Securin (Holt, Krutchinsky et al., 2008) (**Fig. 2B**), more specifically, shifting temperature from 25°C to 28°C shortened the half-life of ^35^S-labeled Securin in extracts from ∼80 to ∼60 minutes. The addition of Ubch10 approximately halved Securin half-life under both conditions (**Fig. 2B**). Temporal dynamics of APC/C^Cdh1^ substrate Tome-1 was vastly different. Similar to Securin, increasing temperature facilitated the electrophoretic mobility shift of Tome-1, known to be associated with mitotic phosphorylation (Ayad et al., 2003, Pe’er, Lahmi et al., 2013). However, Tome-1 remains stable and at a high electrophoretic mobility throughout the experiment (**Fig. 2C**). Importantly, addition of Ubch10 triggered orderly dephosphorylation followed by degradation of Tome-1 in a temperature-controlled manner. These dynamics were evident by the gradual shift of Tome-1 back to its basal electrophoretic mobility form and the subsequent reduction of the ^35^S signal, recapitulating the signaling events of mitotic exit, APC/C^Cdh1^ activation and G1 entry. Thus, at optimal reaction conditions, S3 extracts derived from nocodazole arrested pro-metaphase cells can recapitulate the compete transition from pro-metaphase to G1-like state *in vitro*.

Next, we investigated the behavior of E2F8 in this extracts system. E2F8 degradation was unambiguous only when mitotic exit was accelerated by UbcH10 and commenced considerably after Securin disappearance. The pattern of E2F8 proteolysis was nearly identical to that of Tome-1, suggesting that E2F8 degradation is coordinated with APC/C^Cdh1^ activation (**Fig. 2C-E**). Additionally, the electrophoretic mobility display of E2F8 and Tome-1 shared a remarkable resemblance. This observation suggests an unappreciated regulation of E2F8 by orderly (de)phosphorylation during mitotic progression and exit, as well as highlights the power of ‘dynamic’ mitotic extracts in quantifying phosphorylation and temporal proteolysis of human proteins across cell division landmarks.

### E2F8 degradation by APC/C is Cdh1-specific

Cdc20 *vs.* Cdh1 specificity of APC/C targets is a key element in the overall mechanism underlying orderly cell division in all eukaryotes (Kernan et al., 2018). The kinetics of E2F8 degradation observed above (**Fig. 2D**) are consistent with its regulation by APC/C^Cdh1^, despite prior reports suggesting destruction by both APC/C^Cdc20^ and APC/C^Cdh1^ (Boekhout et al., 2016). To address this discrepancy, we developed a mitotic cell-free system with a constitutively active APC/C^Cdc20^ and inactive APC/C^Cdh1^. High Cdk1 activity prevents Cdh1 from interacting with APC/C (Jaspersen, Charles et al., 1999, Kramer, Scheuringer et al., 2000, Listovsky, Zor et al., 2000, Zachariae, Schwab et al., 1998). Thus, we developed an extract system from mitotic 293-T-REx cells where Cdk1 is constitutively active, due to the tetracycline (tet)-induced expression of a <u>n</u>on-<u>d</u>egradable allele of Cyclin <u>B</u>1 (NDB); hereafter referred to as NDB cells/system (**Fig. 3A**). Non-degradable Cyclin B1 blocks cells in an anaphase-like state, *i.e.,* post-mitotic checkpoint removal (Pe’er et al., 2013, Wheatley, Hinchcliffe et al., 1997, Zur & Brandeis, 2001, Zur & Brandeis, 2002). In the absence of tet, the selected NDB colony tolerates the basal expression of non-degradable Cyclin B1, as evidenced by the normal cell cycle profile and cell size range (**Fig. 3B and C**). Following treatment with tet, virtually all cells arrest in mitosis, displaying a typical round shape and separated sister chromatids (**Fig. 3D-G**). Importantly, APC/C^Cdc20^, but not APC/C^Cdh1^, is active in tet-treated NBD cells. Consequently, Geminin, an APC/C^Cdc20^ substrate (McGarry & Kirschner, 1998)], but not Cdc20, an APC/C^Cdh1^ substrate (Pfleger & Kirschner, 2000), is reduced in tet-induced NDB cells relative to pro-metaphase arrested mitotic cells (**Fig. 3H**). A reduction in Cdc20 could be observed only in G1 cells (**Fig. 3I**). Consistent with the expression of non-degradable Cyclin B1, mitotic extracts derived from tet-induced NDB cells exhibit high Cdk1 activity, highlighted by the electrophoretic mobility shift of Securin and Tome-1 (**Fig. 3J**). Moreover, APC/C^Cdc20^ is highly active, whereas APC/C^Cdh1^ is not, evidenced by the degradation of Securin but not Tome-1 (**Fig. 3J**). As opposed to ‘dynamic’ pro-metaphase extracts made from nocodazole arrested cells (**Fig. 2**), NDB mitotic extracts are ‘*static’* and remain unable to trigger Tome-1 dephosphorylation and degradation by excess of UbcH10 (**Fig. 3K**). Importantly, E2F8 was shifted to a high electrophoretic mobility form following incubation in NDB mitotic extract, further supporting the idea that E2F8 is phosphorylated by mitotic kinases. Like Tome-1, E2F8 was not degraded in NDB extracts. Therefore, experiments in orthogonal cell extract systems, from two different cell lines, suggest that E2F8 is targeted for degradation by APC/C^Cdh1^, but not APC/C^Cdc20^ (**Fig. 3L**). To rule out the possibility that the stability of E2F8 and Tome-1 in extracts derived from HEK293 is cell line specific, we inactivated Cdk1 using the small-molecule inhibitor RO-3306. This treatment successfully overrides the mitotic arrest induced by non-degradable Cyclin B1, resulting in mitotic exit, evidenced by cell morphology, DNA profiling and the canonical dephosphorylated state of the APC/C subunit Cdc27 (Kramer et al., 2000) (**Fig. 3M and N**). Next, we analyzed APC/C complexes by Cdc27 IP in control and RO-3306 treated mitotic NDB extracts. Cdk1 inhibition causes APC/C to dissociate from Cdc20, bind Cdh1, and degrade endogenous Cdc20 (an APC/C^Cdh1^ substrate), recapitulating the canonical APC/C^Cdc20^-to-APC/C^Cdh1^ switch seen *in vivo* during mitotic exit (**Fig. 3O and P).** Furthermore, neither Tome-1 nor E2F8 is mobility-shifted in Cdk1-inhibited extracts (**Fig. 3Q**). Most importantly, E2F8 and Tome-1 were both degraded in RO-3306 treated cell extracts where APC/C^Cdh1^ has been activated. Degradation of both proteins is blocked by dominant negative Ubch10, and is, thus, APC/C dependent (**Fig. 3Q**). We concluded that E2F8 is specifically targeted for degradation by APC/C^Cdh1^.

**Figure 3:**
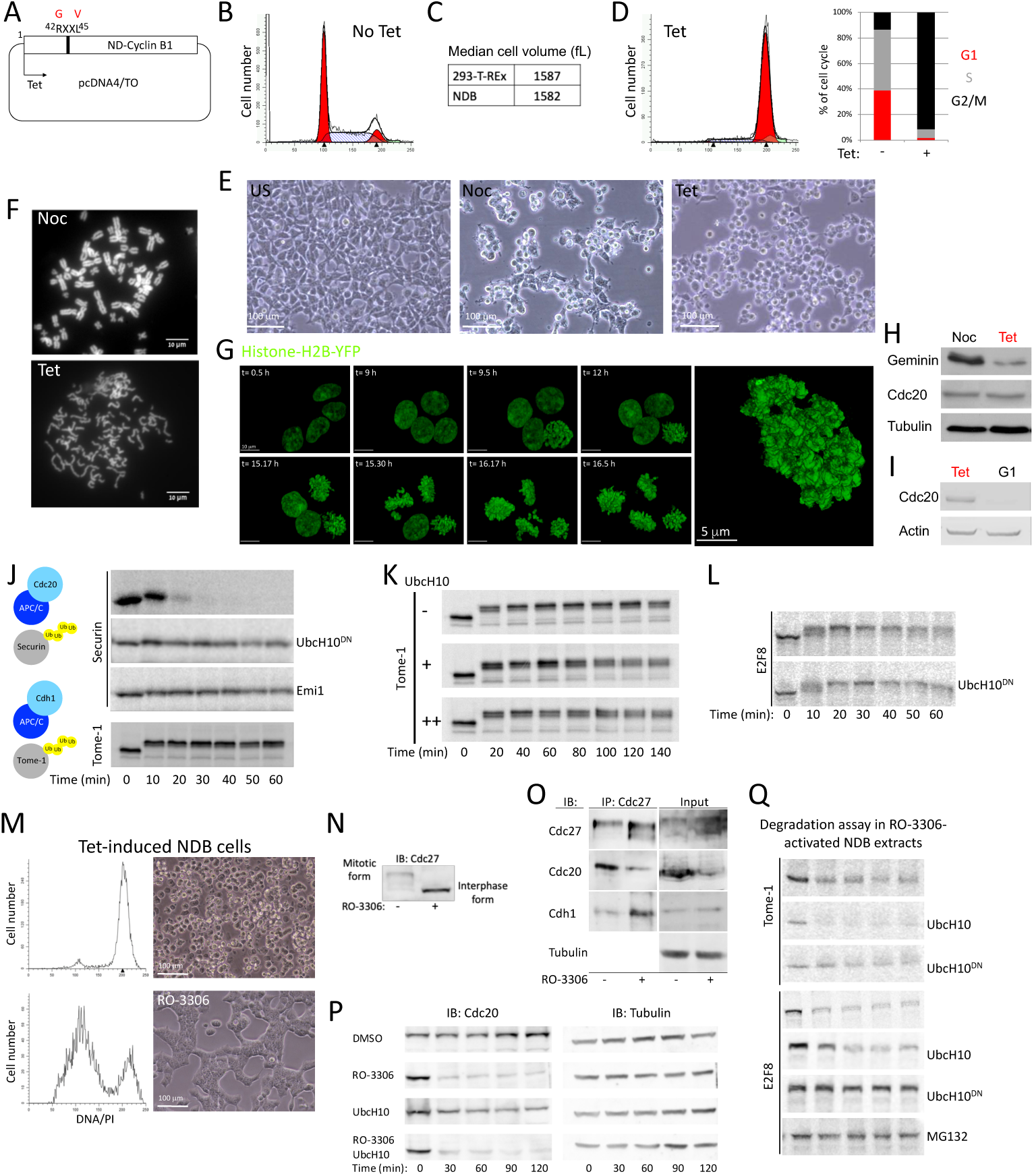
E2F8 is stable in ‘static’ APC/C^Cdc20^-active extracts. **(A)** Human Cyclin <u>B</u>1 in which Arg 42 and Leu 45 are substituted with Gly and Val is <u>n</u>on-<u>d</u>egradable (ND). A colony of 293-T-REx cells stably expressing <u>ND</u>-Cyclin <u>B</u>1 (NDB) under tetracycline (tet)-regulated CMV promoter was generated (*i.e.,* NDB cells). **(B)** DNA distribution of asynchronous NDB cell population. **(C)** Median cell size (fL: femtoliter) of NDB cells *vs.* parental 293-T-REx cells. **(D)** DNA distribution of NDB cells following 22 h treatment with tet. Cell cycle phase distributions of NDB cells pre- and post-induction with tet are shown. **(E)** A panel of light phase images of NDB cells pre- and post-incubation with Nocodazole (Noc) or tet. **(F)** Noc- or tet-treated NDB cells were harvested for chromosome spreads. Representative fluorescent images of DAPI stained chromosomes are shown. **(G)** An image series of live NDB cells stably expressing Histone-H2B following tet treatment. A higher resolution image of a live, tet-treated NDB cell with typical disorganized chromatids is shown on the right. **(H)** Noc- or tet-treated NDB cells were harvested for Western blotting. **(I)** NDB cells showing the lowest 10% forward scatter width (FSC-W) signal were sorted. These G1 cells, as well as tet-treated NDB cells, were harvested for Western blotting with the depicted antibodies. **(J)** Mitotic cell extracts were generated from tet-induced NDB cells. Degradation of Securin and Tome-1 (^35^S-labeled IVT products) was assayed in extracts supplemented with mock, UbcH10^DN^, or APC/C inhibitor Emi1 (C-terminal fragment; recombinant). Time-dependent degradation was assayed by SDS-PAGE and autoradiography. **(K)** Mitotic NDB extracts were supplemented with increasing doses of UbcH10. Degradation and electrophoretic mobility-shift of Tome-1 was examined. **(L)** Degradation assay of E2F8 (^35^S-labeled IVT products) in mitotic NDB extracts supplemented with mock or UbcH10^DN^. **(M-P)** *Tet-induced NDB cells/extracts can exit mitosis into a G1-like state by blocking Cdk1 activity***. (M-N)** DNA distributions, light phase images (M), and immunoblot (IB) (N) of tet-induced NDB cells pre- and post-incubation with RO-3306 (135 min). The mitotic *vs.* G1 electrophoretic mobility-shift of Cdc27 is shown. **(O)** Mitotic extracts were generated from tet-induced NDB cells. Coimmunoprecipitation (IP) of Cdc27 was performed pre- and post-treatment with RO-3306 (30 min). **(P)** Time-dependent dynamics of endogenous Cdc20 in mitotic NDB extracts following incubation with DMSO, RO-3306 and/or UbcH10. Cdc20 levels were detected by Western blotting. Loading control (Tubulin) is also shown. **(Q)** Time-dependent degradation of Tome-1 and E2F8 (^35^S-labeled IVT products) in NDB mitotic extracts pre-activated with RO-3306 (15 min). Reactions were mock treated or supplemented with UbcH10, UbcH10^DN^, or MG132. E2F8 and Tome-1 are stable in APC/C^Cdc20^-active NDB extracts. Both proteins are degraded in extracts where APC/C^Cdh1^ is active.

### E2F8 ubiquitination in G1 is primarily Lys11-linked

Cell extracts generated from synchronous S3 cells 3-3.5 h after release from a nocodazole block exhibit homogenous and optimal APC/C^Cdh1^ activity. These cell extracts, defined here as ‘G1 extracts’, have been used for the discovery of E2F8, E2F7, and other cell cycle proteins as APC/C^Cdh1^ targets (Cohen et al., 2013, Pe’er et al., 2013, Singh, Winter et al., 2014). While Lys(K)48-linked ubiquitination is considered the main signal for proteasomal protein degradation (Kravtsova-Ivantsiv, Sommer et al., 2013), APC/C preference for K11-linked Ubiquitin chains has been demonstrated for an increasing number of substrates, including Securin and Kifc1 (Jin, Williamson et al., 2008, Noach-Hirsh, Nevenzal et al., 2015, Wu, Merbl et al., 2010). To test whether this feature applies to E2F8, we utilized a designated microfluidic platform with which ubiquitination of freshly expressed proteins can be tested in G1 extract (Noach-Hirsh et al., 2015). The assay is based on *in situ* detection of EGFP-tagged substrates (**Fig. S2**) and Rhodamine-labelled Ubiquitin (Rd-Ub) (illustrated in **Figure 4A and B**). Once ubiquitination of E2F8-EGFP was validated on-chip to be APC/C specific (**Fig. 4C**), we tested the displacement of Rd-Ub from E2F8 by 10-fold excess of <u>unlabeled</u> K-to-Arg(R) mutant ubiquitin variants. As expected, excess of unlabeled WT Ubiquitin in the reaction mix outcompeted Rd-Ub, evidenced by the sharp drop of net Rd signal. Similar results were obtained when K48R- or K63R-Ubiquitin mutants (UbK48R, UbK63R) were added. In contrast, the impact of K11R-Ubiquitin mutant (UbK11R) on Rd-Ub signal was mild (**Fig. 4D**), reflecting the low capacity of this particular mutant to form ubiquitin chains on E2F8. Comparable degradation assays in G1 extracts were confirmatory: E2F8 proteolysis became inefficient only when UbK11R was supplemented to the reaction (**Fig. 4E**), suggesting that E2F8 ubiquitination is primarily mediated by K11-linked ubiquitin chains.

**Figure 4:**
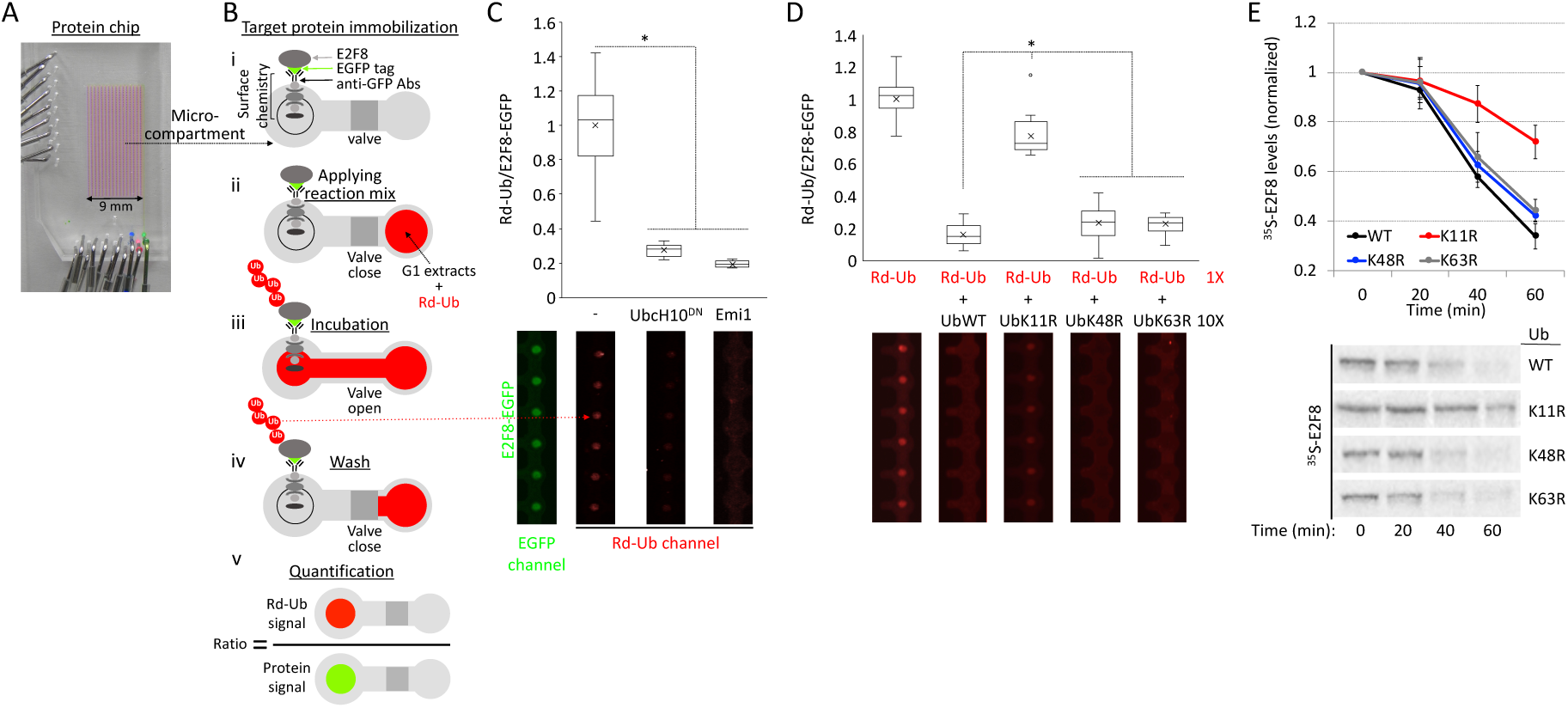
Ubiquitination of E2F8 by APC/C^Cdh1^ is primarily via K11-linked ubiquitin chains. **(A)** An image of an integrated microfluidic platform comprising microcompartments isolated by pneumatic valves. **(B)** Each microcompartment has two chambers. Fresh E2F8-EGFP IVT product, was applied to the chip and immobilized to the ‘Protein chamber’ via anti-GFP antibodies (Abs) and a designated surface chemistry (i). Next, G1 extracts supplemented with Rhodamine-coupled Ubiquitin (Rd-Ub) were applied to the second chamber (ii). The opening of the valve allows reaction mix to diffuse into protein chambers, allowing ubiquitination of the immobilized substrate (iii). After 10 min incubation, all protein chambers are washed (iv). Rd-Ub moieties attached to E2F8-EGFP at the protein chamber are quantified by a fluorescence imaging. Rd-Ub signal in each protein chamber is normalized to E2F8-EGFP levels, *i.e.,* ‘Protein signal’ (v). **(C)**. *APC/C^Cdh1^-mediated ubiquitination of E2F8 on-chip*. E2F8-EGFP was expressed in reticulocyte lysate, deposited on the chip surface, and incubated with G1 extracts supplemented with either mock, UbcH10^DN^ or Emi1. Normalized Rd-Ub signals were calculated from 20 microcompartments (mean [X], median [–], and 4 quantiles (box and whiskers) are indicated; \**p* value < 0.001). Array sections showing ‘raw’ Rd-Ub signals of six microreactions for each of the three conditions are shown (red dots). A representative image of immobilized E2F8-EGFP is also shown (green dots). **(D)** Ubiquitination of E2F8-EGFP was assayed in the presence of G1 extracts, Rd-Ub, and excess of <u>unlabeled</u> WT or mutant Ubiquitin in which Lys 11 (UbK11R), Lys 48 (UbK48R), or Lys (UbK63R) were substituted with Arg. Plots average 18 microreactions. Array sections showing ‘raw’ Rd-Ub signals are depicted. **(E)** Degradation of ^35^S-labeled E2F8 (IVT product) was assayed in G1 extracts supplemented with WT or mutant Ubiquitin. Time-dependent degradation was assayed by SDS-PAGE and autoradiography. Mean and SE values are plotted (*n* = 3). ^35^S-E2F8 signals are normalized to *t* = 0. A set of source data is shown.

### Multiple functional motifs coordinate E2F8 proteolysis in G1

Direct assays in G1 extracts have been proven informative in mapping and characterizing destruction motifs of APC/C substrates (Jin et al., 2008, Pe’er et al., 2013, Singh et al., 2014, Wu et al., 2010). E2F8 carries multiple motifs potentially associated with APC/C-mediated degradation– three KEN sequences and two RXXL sequences (**Fig. 5A**). We generated point-mutations in each of these motifs (**Fig. 5B**) and quantified the kinetics of E2F8 proteolysis in G1 extracts. RXXL-to-GXXV mutation at position 183 inhibited E2F8 degradation (**Fig. 5C**). Mutations in each of the previously reported KEN boxes at position 5 and 375 (Boekhout et al., 2016) had a borderline impact on E2F8 proteolysis. However, the two KEN box mutations had a noticeable impact on E2F8 degradation when combined with mutant RXXL 183 or with each other (**Fig. 5D**). Mutations in RXXL 87 and KEN 637 had no inhibitory effect (**Fig. 5C and D**). The functionality of KEN 5 and KEN 637 was further assayed in the context of short N- and C-terminal fragments [(E2F8-N80 and E2F8-C (**Fig. 5E**)], reasoning that this approach could highlight the potential potency of single elements in driving E2F8 degradation. The E2F8 C-terminus was stable, consistent with this region not regulating proteolysis. Instead, degradation of E2F8-N80 was efficient and KEN specific (**Fig. 5F**). Overall, we conclude that RXXL 183, KEN 5, and KEN 375, but not RXXL 85 and KEN 657 mediate E2F8 degradation by APC/C^Cdh1^. This result also highlights the ability of multiple degrons to contribute cooperatively to substrate proteolysis, perhaps through multivalent APC/C binding (Watson, Brown et al., 2019).

**Figure 5:**
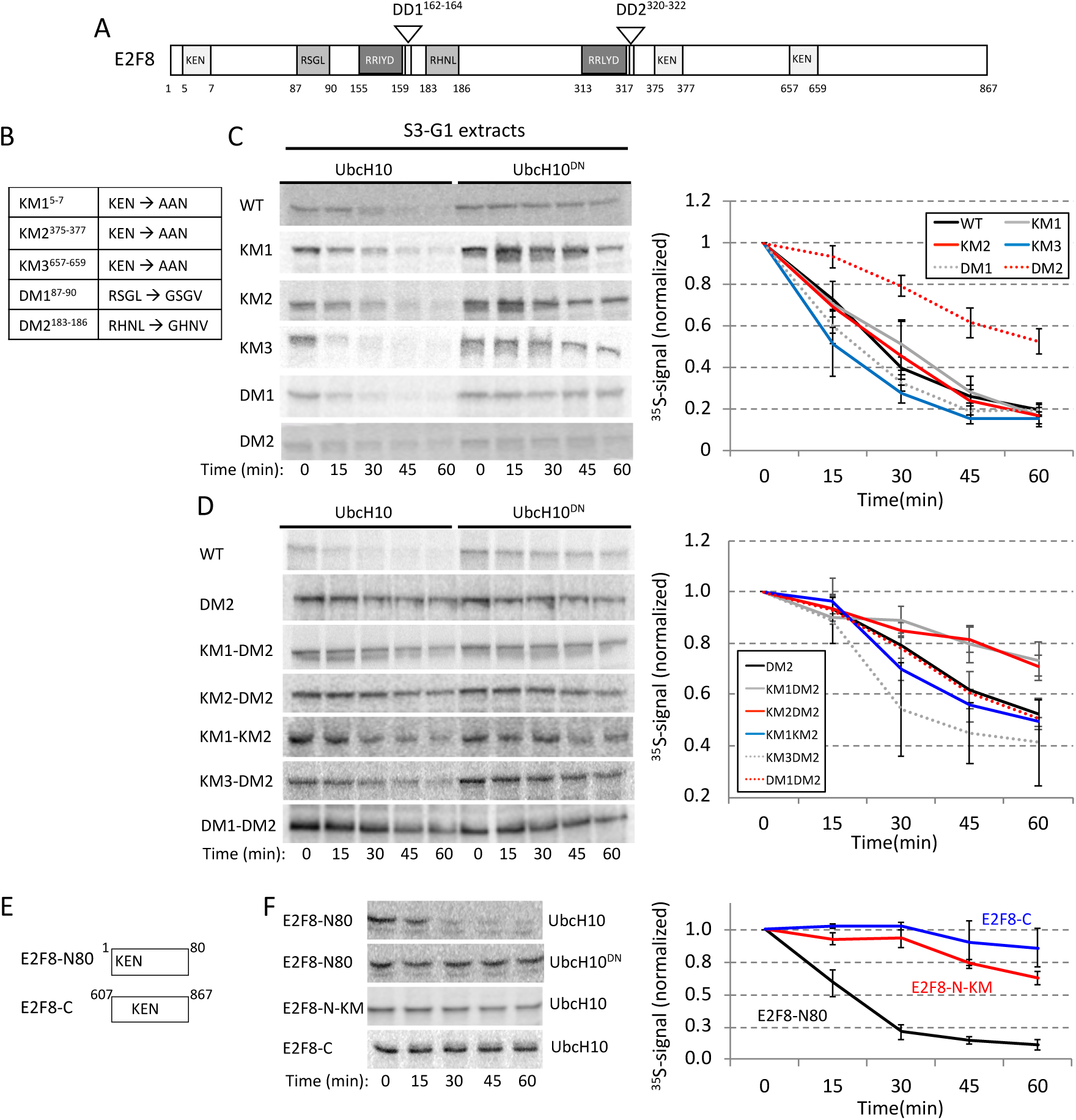
Multiple functional motifs coordinate E2F8 proteolysis in G1. **(A)** Schematics of human E2F8. KEN and RXXL motifs are shown alongside the conserved DNA-binding (RRXYD)- and <u>d</u>imerization (DD1, DD2) <u>d</u>omains. **(B)** A list of E2F8 mutant variants generated by site-directed mutagenesis. Amino acid substitutions are indicated for each of the five KEN/RXXL motifs. KM1/2/3: <u>K</u>EN <u>m</u>utant <u>1/2/3</u>; DM1/2: <u>D</u>estruction-box <u>m</u>utant <u>1/2</u>. **(C)** Degradation of ^35^S-labeled E2F8 variants (IVT products) was tested in G1 extracts supplemented with UbcH10 or UbcH10^DN^. Time-dependent degradation was assayed by SDS-PAGE and autoradiography. Representative raw data and quantifications are shown. Mean E2F8 levels (^35^S signals) normalized to max signal at *t* = 0 are shown (*n* = 3-4). Bars represent SE **(D)**. E2F8 double mutants were analyzed as described in C. **(E)** Schematics of N- and C-terminal fragments of E2F8 (E2F8-N80/C) carrying a single KEN motif. **(F)** Time-dependent degradation of E2F8 fragments (see details in C). Overall, the data suggest that RXXL^183^, KEN^5^, and KEN^375^, but not RXXL^85^ and KEN^657^, contribute cooperatively to E2F8 degradation by APC/C^Cdh1^.

### E2F8 proteolysis in G1 is mediated by N-terminal Cdk1 Sites

The temporal electrophoretic mobility of E2F8 in ‘dynamic’ (**Fig. 2**) and ‘static’ (**Fig. 3**) mitotic extracts can be explained by orderly phosphorylation and dephosphorylation during mitotic progression and exit. There are four T/SP sites in E2F8-N80 fragment, two of which are TPxK, *i.e*., the canonical Cdk1 consensus sites (**Fig. 6A**). Mobility shift of both full-length E2F8 and N80-E2F8 in NDB mitotic extracts was blocked by Cdk1 inhibitor (**Fig. 6B**). Thr(T)-to-Ala(A) mutation in position 20 or 44 reduced the mobility shift of E2F8-N80, and all the more so when combined (**Fig. 6C**). We concluded that Cdk1/Cyclin B1 phosphorylates E2F8 in mitosis at position T20 and T44. Phosphorylation in proximity to destruction motifs can regulate APC/C-mediated ubiquitination (Holt et al., 2008, Singh et al., 2014). This mechanism helps coordinate degradation with the cell cycle clock (Holt et al., 2008). We, therefore, tested the potential link between E2F8 phosphorylation and degradation. The E2F8-N80 fragment (**Fig. 6C**) allowed us to focus on the potential relationship between T20/T44 phosphorylation and KEN box at position 5. Alanine mutations at T20 and/or T44 had no impact on the degradation of E2F8-N80 in G1 extracts (**Fig. 6D**). In contrast, phosphomimetic mutation [T-to Asp(D)] at T44 partially impeded E2F8-N80 degradation. This inhibitory effect was vastly increased when both T20 and T44 were substituted with D (**Fig. 6D**). Most importantly, T20D/T44D mutation strongly impaired the proteolysis of full-length E2F8 in G1 extracts (**Fig. 6C**). In fact, the impact of T20D/T44D mutation on E2F8 half-life in G1 extracts was greater than any of the single KEN or RXXL mutations (**Fig. 5C**). Altogether, our findings couple E2F8 ubiquitination by APC/C^Cdh1^ with an unphosphorylated state of T20 and T44. This molecular switch can restrict E2F8 degradation by APC/C to times of low Cdk1 activity.

**Figure 6:**
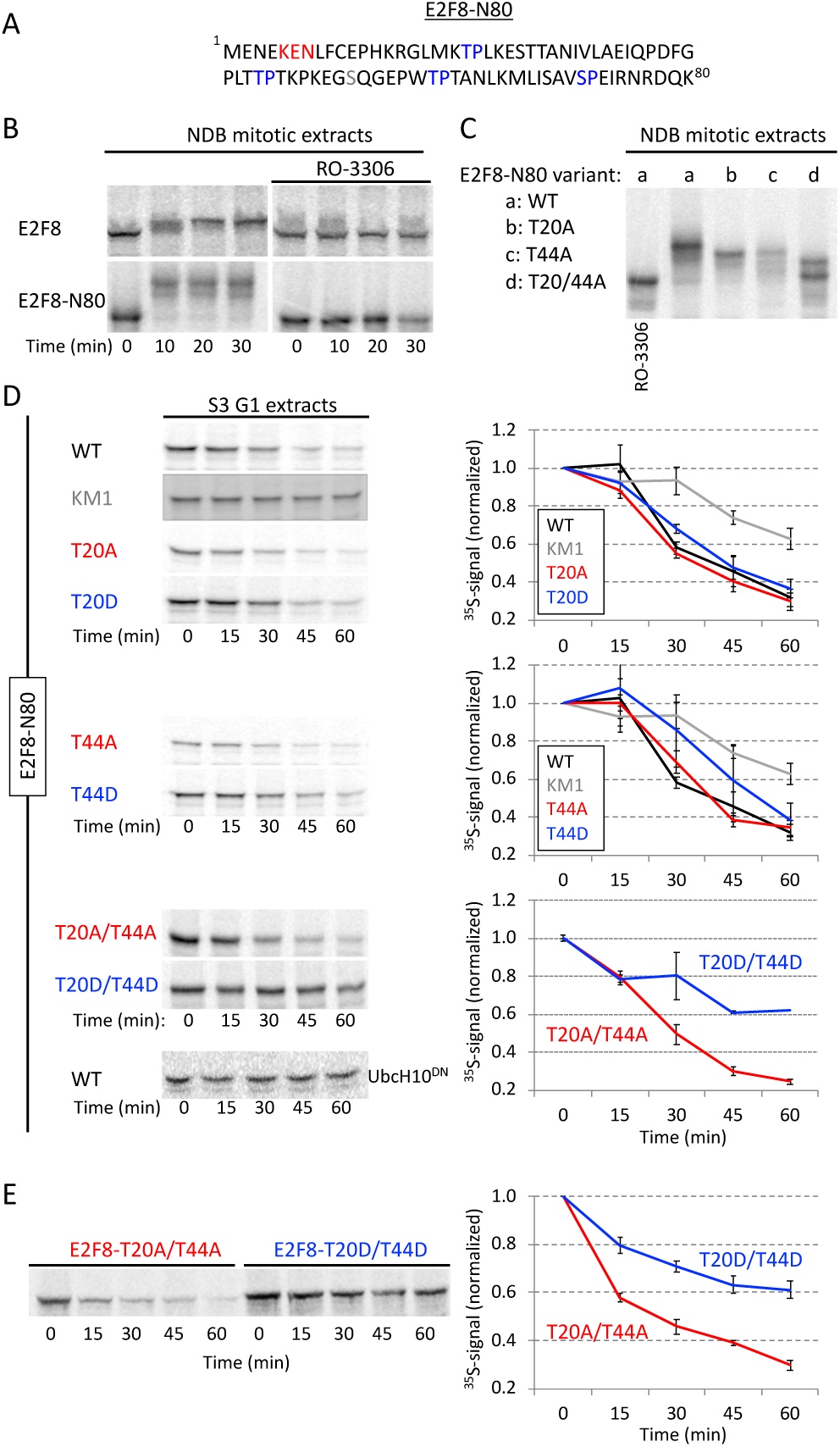
Phosphomimetic Cdk1 sites stabilized E2F8 in G1 extracts. **(A)** E2F8 N-terminal fragment of 80 amino acids (E2F8-N80). KEN box and 4 canonical Cdk1 consensus phosphorylation sites are colored. **(B)** Time-dependent electrophoretic mobility shift of full-length and E2F8-N80 (^35^S-labeled IVT products) in NDB mitotic extracts supplemented with mock or the Cdk1 inhibitor RO-3306. **(C)** Thr (T)20 and/or T44 of E2F8-N80 were substituted with Ala (A). Mobility shifts of WT *vs.* mutant E2F8-N80 variants are shown. **(D)** Time-dependent degradation in G1 extracts of E2F8-N80 and the following variants: KEN box mutant (KM1), single/double phosphomimetic mutants [T-to-Asp (D)], and single/double phospho-dead mutants (T-to-Ala). **(E)** Time-dependent degradation of full-length E2F8 carrying double phosphomimetic or double phospho-dead mutations. (B-E) Protein degradations and electrophoretic mobility-shifts were assayed by SDS-PAGE and autoradiography. A set of source data is shown. Mean and SE calculated from three degradation assays are plotted.

### Temporal proteolysis of E2F8 across the cell cycle

The contribution of temporal proteolysis to the overall dynamics of E2F8 across the cell cycle (**Fig. 1**) has yet to be addressed systematically. To this end, seven cell-free systems were generated, recapitulating cell cycle milestones from pro-metaphase to late S-phase. Degradation of E2F8 was assayed alongside Securin and p27/Kip1. The latter was used as a positive control for SCF^Skp2^ activity (Carrano, Eytan et al., 1999). To minimize non-physiological variabilities, protein concentrations of all extracts were matched (+/− 10%), only Ubiquitin and energy-regeneration mixture were supplemented to the reactions, and all assays were performed at 28°. The stability of E2F8 and delayed proteolysis of Securin in pro-metaphase extracts (Pro-M) were both expected at 28°C (**Fig. 7A and Fig. 2B and D**). The stability of E2F8 in G1-S extracts and in all the three S-phase extracts was in agreement with the temporal dynamics of the protein at these stages (**Fig. 7A and Fig. 1**). p27 proteolysis validated the specific activity of mid- and late S-phase extracts. Proteolysis of both E2F8 and Securin in early-mid G1 extracts was optimal, reflecting the high APC/C^Cdh1^ activity and lowest abundance of E2F8 associated with this stage (**Fig. 7A**). Interestingly, the degradation of Securin was slightly more efficient than E2F8 (**Fig. 7A and B**), potentially reflecting the high ubiquitination efficiency of the former protein (Rape et al., 2006). More importantly, APC/C activity has been shown to weaken during late G1 (Huang, Park et al., 2001, Rape & Kirschner, 2004). Accordingly, the half-life of Securin was indeed slightly longer in late G1 extracts, prepared 6 hours after the nocodazole release. However, while the overall degradation of Securin remained potent in late G1 extracts, E2F8 proteolysis nearly ceased (**Fig. 7A and B**). The differential stability of E2F8 *vs.* Securin in late G1 extracts effectively demonstrates an intrinsic mechanism by which E2F8 can accumulate while APC/C is still active, providing a mechanism for E2F8 buildup during late G1 (**Fig. 1A**).

**Figure 7:**
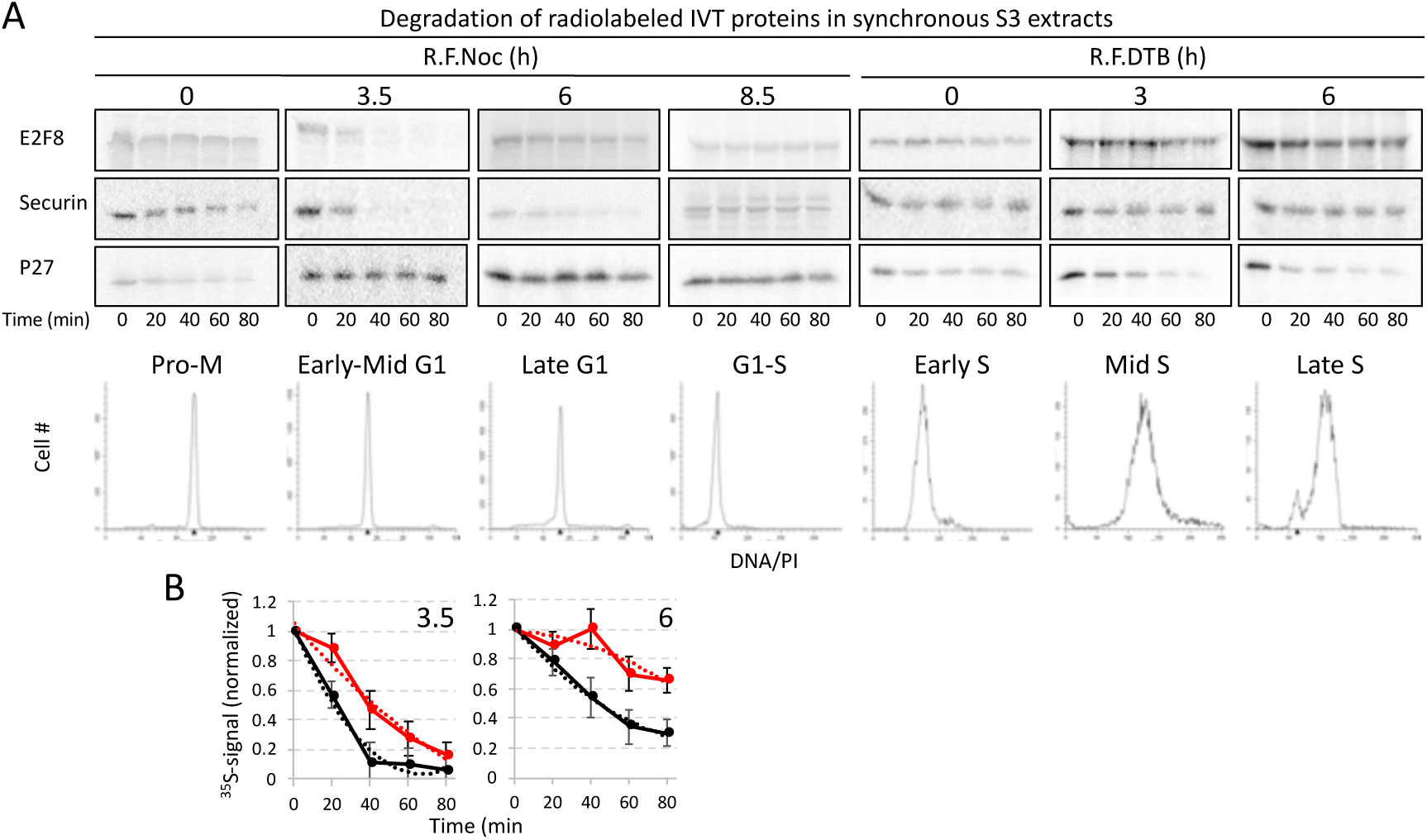
*Temporal proteolysis of E2F8* across the cell cycle. **(A)** Degradations of ^35^S-labeled E2F8, Securin and p27 (IVT products) were tested in seven cell extracts generated from synchronous S3 cells at seven points across the cell cycle (R.F.Noc: Release from a thymidine-nocodazole block; R.F.DTB: Release from a double thymidine block). DNA distributions are shown. Extracts were supplemented only with ubiquitin and energy-regeneration mix. Time-dependent degradation was assayed by SDS-PAGE and autoradiography. A set of source data is shown. **(B)** Average, SE and polynomial fit (dotted line) calculated from three degradation assays of E2F8 and Securin in early-mid- and late-G1 extracts. ^35^S signals were normalized to *t*_0_. The differential degradation capacity for E2F8 *vs.* Securin in late G1 extracts is demonstrated.

### E2F8 is regulated by SCF^Cyclin F^ E3 ligase

The reduction in E2F8 levels seen pre-mitosis (**Fig. 1**) is likely to be regulated at both transcriptional (E2F1) and post-translational levels. The SCF-family of ubiquitin ligases represent attractive candidates for mediating E2F8 destruction owing to their key roles in the cell cycle (Silverman, Skaar et al., 2012). SCF ligases utilize substrate adaptor F-box proteins that bind specific substrates and mediate their ubiquitination and degradation. SCF^Skp2^, SCF^β-TrCP^, and SCF^CyclinF^ have all been implicated in cell cycle and are candidates for driving E2F8 downregulation pre-mitosis although none have been linked to atypical E2Fs. Among those three, several pieces of evidence nominate Cyclin F as the most likely candidate. First, Skp2 is active when E2F8 levels peaks, and E2F8 is stable in S-phase extracts that recapitulate p27 degradation, a well-established SCF^Skp2^ substrate (**Fig. 7A**). Second, SCF^β-TrCP^ binds to a well-defined recognition sequence (DSGXX(XX)S) that is absent in E2F8 (Orian, Gonen et al., 2000, Yaron, Hatzubai et al., 1998). Third, Cyclin F peaks at G2 phase (Choudhury, Bonacci et al., 2016, D’Angiolella, Esencay et al., 2013, Galper, Rayner et al., 2017, Mavrommati, Faedda et al., 2018), concomitantly with E2F8 downregulation (**Fig. 1**). And fourth, E2F8 carries putative RxL motifs (Cy box) (**Fig. 8A**) that are known to mediate ubiquitination by SCF^CyclinF^. We were unable to reproduce SCF^Cyclin F^ activity in cell extracts, and such a system has yet to be reported by others. Therefore, we examined the functional relationship between E2F8 and Cyclin F *in vivo*.

E2F8 levels were elevated in cells where Cyclin F has been <u>k</u>nocked <u>o</u>ut (KO) using CRISPR-Cas9 (**Fig. 8B**). In addition, Cyclin F overexpression reduced the levels of ectopically expressed E2F8, suggesting that Cyclin F promotes E2F8 degradation (**Fig. 8C**). To analyze endogenous E2F8, we constructed cells with doxycycline-inducible Cyclin F expression in MCF7 and T47D cell lines. In both, Cyclin F expression caused a dose-dependent decrease in E2F8 abundance (**Fig. 8D**). Together, this suggests that Cyclin F regulates E2F8 degradation via ubiquitin mediated proteolysis. Accordingly, treatment with proteasome inhibitors MG132 and bortezomib prevented the degradation of endogenous E2F8 in MCF7 cells expressing doxycycline-inducible Cyclin F (**Fig. 8E**). We conclude that Cyclin F promotes the degradation of E2F8 through the proteasome. Next, we examined E2F8 abundance in control and Cyclin F KO cells traversing G2. Cells were released from synchronization at G1/S by double thymidine block and analyzed by immunoblotting. A reduction in E2F8 was evident in late S and G2-phase, prior to mitotic entry, in control cells (**Fig 8F**). However, E2F8 levels persisted into mitosis in Cyclin F KO cells, strongly suggesting that the degradation of E2F8 observed in G2 is Cyclin F dependent. Owing to the role that RxL motifs often play in SCF^CyclinF^-mediated ubiquitination, we analyzed E2F8 variants in which the Arginine of each of the four RxL motifs was substituted with Alanine (RxL-to-AxL). Similar to the result above, Cyclin F overexpression strongly downregulated the abundance of wild-type, exogenously expressed E2F8 (**Fig. 8G**). However, the mutation of R313 resulted in limited sensitivity to Cyclin F overexpression, whereas the three other E2F8 alleles (R15A, R81A and R587A) behaved identically to the wild-type protein (**Fig. 8G**). This suggests that the regulation of E2F8 by Cyclin F is mediated, at least in part, by the RxL motif at position R313. It is noteworthy that RRL 313 is part of E2F8 DNA binding domain (**Fig. 5A**) and the expression of R313 mutant was lower compared to all other mutated variants (**Fig. 8F**). While the possible impact of an R313A mutation on the overall structure of E2F8 cannot be ignored, E2F8 carrying mutation in both DNA binding domains was degraded by APC/C^Cdh1^ (**Fig. S3**). We next analyzed the possibility that E2F8 and Cyclin F interact. We expressed FLAG-Cyclin F and 6HIS-E2F8-HA in HEK293T cells and analyzed their interaction by co-IP. We detected an interaction between the two proteins, irrespective of which protein was precipitated (**Fig. 8H and 8I**), suggesting that E2F8 binds Cyclin F. Taken together, these data strongly suggest that E2F8 regulation by Cyclin F is direct and that SCF^Cyclin F^ activity downregulates E2F8 prior to M-phase entry. These data complement the recent discovery that E2F1 and other members of the activating branch of the E2F family are targeted for proteasomal degradation by the SCF^Cyclin F^ complex (Clijsters, Hoencamp et al., 2019).

**Figure 8:**
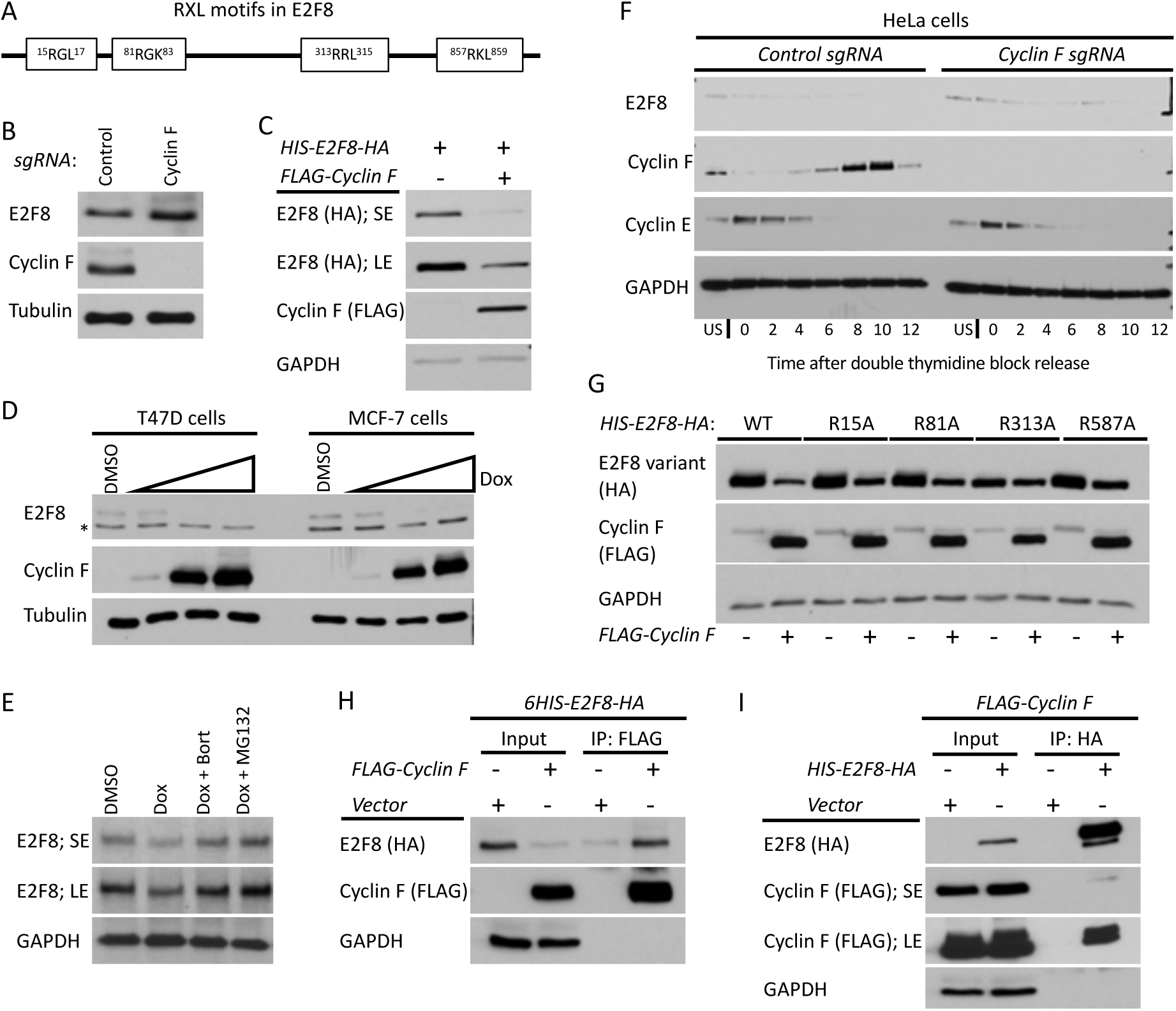
Cyclin F mediates the degradation of E2F8 in G2-phase. **(A)** Schematic depiction of the RxL motifs in E2F8. **(B)** The abundance of E2F8 in control Cyclin F KO HeLa cells was analyzed by western blot. **(C)** HEK293T cells were transiently transfected with HIS-E2F8-HA with and without FLAG-Cyclin F. Cells were analyzed by western blot 48 hours post transfection. SE: short exposure; LE: long exposure. **(D)** MCF7 and T47D were engineered to express doxycycline (Dox) inducible Cyclin F. Cells were treated with doxycycline at increasing concentrations for 48 hours and then endogenous E2F8 was analyzed by western blot. **(E)** MCF7 cells were treated with doxycycline to induce expression of Cyclin F. Eight hours prior to harvesting for western blot, cells were treated 8h with either of two proteasome inhibitors, MG132 and bortezomib. Endogenous E2F8 was analyzed by western blot. **(F)** Control and Cyclin F KO HeLa cells were synchronized at the G1-S boundary by a double thymidine block. Following release from the second thymidine block, samples were collected for western blot analysis at the indicated time points. **(G)** HEK293T cells were transfect with wild-type or mutant versions of HIS-E2F8-HA harboring alanine substitutions at the indicated RxL motifs shown in (A). Their response to ectopic co-expression was analyzed by western blot 48 hours after transfection. **(H and I)** HIS-E2F8-HA and FLAG-Cyclin F were co-transfected into HEK293T cells. Cell lysates were subjected to co-IP with wither anti-HA or anti-FLAG antibodies.

## Discussion

In this study, we developed and utilized human cell-free systems, faithfully recapitulating the post-translational signaling that underlies the core of the cell cycle oscillator, most notably, the phosphorylation and ubiquitination events that promote human cell cycle progression. We discovered multiple mechanisms underlying temporal proteolysis of E2F8 in proliferating cells, with relevance to the overall coregulation between E2F8, E2F1 and the cell cycle clock.

Dynamic mitotic extracts were highly informative, revealing orderly phosphorylation, dephosphorylation and proteolysis of E2F8 while traversing spindle checkpoint inactivation, the APC/C^Cdc20^-to-APC/C^Cdh1^ switch, and G1 entry (**Fig. 2**). To the best of our knowledge, a complete transition of cell extracts from pro-metaphase to G1 has not been demonstrated in somatic cell systems. A complementary system generated from anaphase-like cells (NDB) allowed us to investigate E2F8 under ‘static’ environment of high Cdk1 and APC/C^Cdc20^ activity (**Fig. 3**). Using these two cell-free systems we i) identified T20 and T44 as Cdk1 phosphorylation sites (**Fig. 6**); ii) demonstrated that E2F8 is regulated specifically by APC/C^Cdh1^, and not APC/C^Cdc20^ (**Fig. 3**); and iii) coupled temporal proteolysis of E2F8 with mitotic exit dephosphorylation and APC/C^Cdh1^ activation (**Fig. 2**). In the context of Cdc20-specific activity, NDB extracts are particularly advantageous. First, this system obviates dependency on *in vivo* assays in which Cdc20, an essential protein with a substantial impact on the cell cycle, is either overexpressed or knocked-down for dozens of hours. Second, it enabled us to demonstrate that E2F8, although low-leveled, is, in fact, stable in pro-metaphase and early mitosis (**Figs. 1 and 3**). This uncommon phenomenon is difficult to detect *in vivo*, let alone for a protein whose half-life during both G2 and G1 is short (**Figs. 1, 7 and 8**).

Using G1 extracts, we found E2F8 ubiquitination to be primarily K11-linked (**Fig. 4**), mapped three destruction motifs in E2F8 and ranked their individual and cooperative contribution to APC/C^Cdh1^-mediated proteolysis (**Fig. 5**). Importantly, we discovered that phosphomimetic E2F8 is stable in G1 (**Fig. 6**). This finding, together with the dephosphorylation dynamics revealed prior to E2F8 proteolysis during mitotic exit (**Fig. 2**), suggests that dephosphorylation of E2F8 is a necessary pre-requisite for its degradation by APC/C^Cdh1^. That S/T20 and T44 Cdk1 phosphorylation sites are conserved in atypical E2Fs across vertebrates, further suggests that this molecular switch is fundamental and likely applies to E2F7. Experiments in G1 extracts also illuminated an unexpected role for dimerization domains in E2F8 proteolysis (**Fig. S3**). Whether these results genuinely couple E2F8 dimerization with degradation or, alternatively, mirror a global structural change with implications on folding, function, and regulation of E2F8, await further investigation (see **Figure S3** for more details).

The half-life of E2F8 in G1 was analyzed in two distinct points (**Fig. 7**). In early-mid G1 extracts, E2F8 proteolysis was optimal, although slightly less efficient than Securin. In late G1, however, E2F8 proteolysis nearly ceased while the overall degradation of Securin stayed potent (**Fig. 7B**). These dynamics demonstrate that E2F8, unlike Securin and probably other APC/C substrates, is differentially stabilized during late G1, concomitantly with the declining activity of APC/C (Meyer & Rape, 2011, Rape & Kirschner, 2004). Because E2F1 is already active at this stage, this feature, by itself, allows E2F8 to accumulate and potentially coregulate S-phase entry while APC/C^Cdh1^ is still active (**Fig. 1**). Enhanced stability under sub-optimal APC/C activity is a feature of ‘distributive’ APC/C^Cdh1^ targets, *i.e.,* substrates that must associate with the APC/C multiple times to obtain a proteolytic ubiquitin chain, *e.g.*, Cyclin A. (Rape et al., 2006). Consequently, distributive substrates are the first to become stable under limited APC/C activity because of i) the higher chance of deubiquitinating isopeptidases to strip the emerging ubiquitin chains; and ii) the competition with ‘processive’ substrates like Securin, which undergo multi-ubiquitination in single binding event (Meyer & Rape, 2011). Interestingly, both E2F8 and Cyclin A are E2F1 target regulating the G1-S transition. The early rise of E2F1 in G1 is hard to reconcile with previous *in vivo* studies, suggesting that E2F1 is an APC/C^Cdh1^ target (Budhavarapu, White et al., 2012). We further note that E2F1, unlike E2F8 and E2F7, is stable in G1 extracts (**Fig. S1**).

E2F8 was stable in all three S-phase extracts, in accord with the high levels of the protein throughout S-phase (**Fig. 1**). These results suggest that E2F8 dynamics in continuously proliferating cells is less likely to be regulated by SCF^Skp2^-mediated ubiquitination (**Fig. 7A**). Both E2F1 and E2F8 levels diminished before pro-metaphase (**Fig. 1**). This temporal dynamic is rare among the more than 100 known APC/C targets (Meyer & Rape, 2011). While SCF^Cyclin F^-mediated proteolysis accounts for E2F1 reduction during late-S and G2 phases (Clijsters et al., 2019), how atypical E2Fs are regulated in G2 was unknown. Here, we show that Cyclin F interacts and downregulates E2F8 in a proteasome dependent manner (**Fig. 8**), implying a direct link between E2F8 and SCF^Cyclin F^-mediated ubiquitination. The elevated levels of E2F8 observed in Cyclin F-KO cells during G2, where Cyclin F protein peaks, further emphasize the capacity of SCF^Cyclin F^ to downregulate E2F8 pre-mitosis.

Altogether, our findings support the model depicted in Figure 9. SCF^Cyclin F^ downregulates E2F8 in G2, following E2F1 reduction. Since Cyclin F is degraded in mitosis, low-leveled E2F8 remains stable during early mitosis when APC/C^Cdc20^ is active. Conceptually, this mechanism can allow E2F8 to block residual, unwanted E2F1 activity before cell division is successfully terminated, and is consistent with overactivation of E2F1 having negative consequences on chromosome segregation fidelity (Manning, Longworth et al., 2010, Pfister, Pipka et al., 2018). Cdk1 phosphorylates E2F8 on Thr 20 and Thr 44 during mitotic entry. This module has a stabilizing effect on E2F8. Cdk1 activity dropped during mitotic exit. Subsequent dephosphorylation of both E2F8 and Cdh1 triggers E2F8 degradation by APC/C^Cdh1^. E2F1 accumulates in early-mid G1 under high APC/C^Cdh1^ activity, inducing its own expression alongside expression of E2F8, E2F7 and other target genes that promote S-phase entry (*e.g*., Cdc6, Cyclin E). As long as E2F7 and E2F8 are destabilized by APC/C^Cdh1^, E2F1 activity is unrestrained. Maximizing transcription capacity at this stage might be critical for low-leveled E2F1 to ignite the positive feedback required for its autocatalytic increase that drives cells into S-phase (Johnson, Ohtani et al., 1994). E2F8 is differentially stabilized under limited APC/C^Cdh1^ activity associated with late G1. This feature allows E2F8 to accumulate during late G1, while APC/C^Cdh1^ is still active and mediate proteolysis of Securin and probably other substrates. Because E2F1 levels are already high, the E2F1-E2F8 negative feedback can be formed already in G1 to balance the transcriptional activity of E2F1. Overall, these inter-dynamics ensure a safe transition into S-phase (**Fig. 9**). In this study we focused on E2F8. However, owing to the overall similarity of atypical E2Fs, it is easy to speculate that the model herein applied to E2F7 as well.

**Figure 9:**
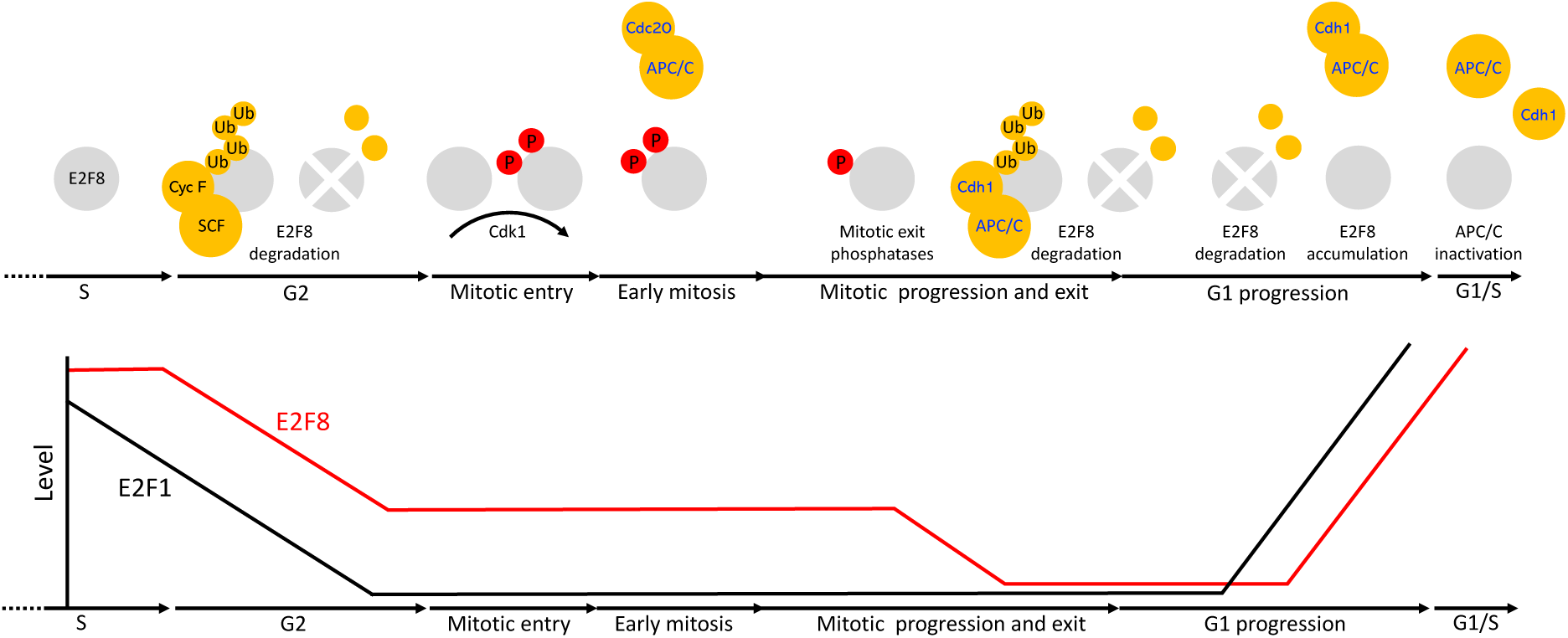
Multiple mechanisms coordinate the dynamics of E2F8 in cycling mammalian cells. **(A)** At the transcriptional level, E2F8 is primarily regulated by E2F1 via a negative feedback mechanism. Post-translationally, E2F8 is controlled by temporal proteolysis orchestrated by multiple pathways. E2F8 peaks in S-phase. During G2-phase, E2F8 protein is downregulated by SCF^Cyclin F^ activity. Although low-leveled, E2F8 remains stable during early mitosis while APC/C^Cdc20^ is active. E2F8 is phosphorylated in mitosis by Cdk1. This phosphorylation has a stabilizing effect on the protein. During mitotic exit, Cdk1 is inactivated and both E2F8 and Cdh1 are dephosphorylated. This dual molecular switch initiates both the assembly of APC/C^Cdh1^ and its ability to ubiquitinate E2F8. The levels of E2F8 remain minimal through G1 as long as APC/C^Cdh1^ is fully active. During late G1, APC/C^Cdh1^ activity weakens by an autonomous mechanism. E2F8 enhanced sensitivity to suboptimal APC/C^Cdh1^ activity effectively stabilizes the protein while other APC/C targets are still degraded. Because E2F1 is already present, the negative feedback circuitry between E2F1 and E2F8 can be formed already in G1 in ensuring a safe transition into S-phase.

## Acknowledgements

We thank Tamar Listovsky-Yulzary, Doron Ginsberg, Vamsi Mootha and Yaron Shav-Tal for sharing reagents. The Tzur lab is supported by the Israel Cancer Research Fund (ICRF), Grant no. RCDA00102, and the Israel Science Foundation (ISF) Grant no. 659/16. The Emanuele lab is supported by the UNC University Cancer Research Fund, National Institutes of Health (R01GM120309), American Cancer Society (RSG-18-220-01-TBG) and donations from the Brookside Foundation.

## Author contributions

D.W., S.N., M.C., T.P.E., M.N.H., J.P., S.Z.P., N.A., E.C., N.L.D., H.N., D.G., M.J.E., and A.T. designed the research.

D.W., S.N., M.C., T.P.E., M.N.H., J.P., S.Z.P., N.A., E.C., N.L.D., X.W., H.N., and R.L. performed the research.

D.W., S.N., M.C., T.P.E., M.N.H., J.P., S.Z.P., N.A., E.C., D.G., M.J.E., and A.T. contributed new reagents or analytic tools.

D.W., S.N., M.C., T.P.E., M.N.H., J.P., S.Z.P., N.A., E.C., N.L.D., X.W., H.N., and R.L., analysed the data.

D.W., S.N., M.N.H., M.J.E., and A.T. wrote the paper.

## Material and Methods

### Plasmids

The following plasmids were used in previous studies: 1) pCS2-FA-E2F8 (Cohen et al., 2013); 2) pCS2-FA-E2F7 (Cohen et al., 2013); 3) pCS2-FA-Securin (Pe’er et al., 2013); 4) pCS2-Tome-1 (*Xenopus laevis*) (Ayad et al., 2003); 5) FLAG-Cyclin F (Choudhury et al., 2016). Histone H2B-YFP plasmid was a gift from Yaron Shav-Tal (Bar-Ilan University). The following plasmids were generated for this study: 1) pCS2-FA-E2F8-EGFP; E2F8 open reading frame (ORF) was amplified by <u>p</u>olymerase <u>c</u>hain <u>r</u>eaction (PCR) using pCS2-FA-E2F8 as a template and primers flanked with FseI (forward) and AgeI (reverse) restriction enzyme (RE) sites, and cloned into pCS2-FA-EGFP vector; 2) pCS2-FA-P27; ORF of p27 was amplified from cDNA template (Open Biosystems) and cloned into pCS2-FA vector using FseI (5’) and AscI (3’) RE; 3) pCS2-FA-His-E2F8-HA; E2F8 ORF was amplified with a forward primer flanked with FseI RE site and a reverse primer flanked with HA tag followed by AscI RE site. The PCR product was cloned downstream to His tag; 4) pCS2-His-E2F8-N80-EGFP; The ORF encoding for amino acid 1-80 of E2F8 was amplified using pCS2-FA-E2F8 template, and a primer set flanked with FseI (forward) and AgeI (reverse) RE sites. The PCR product was cloned into pCS2-FA-EGFP vector downstream to His tag and upstream to EGFP; 5) pCS2-E2F8-C; The ORF encoding for amino acid 607-867 of E2F8 were amplified using pCS2-FA-E2F8 template and a primer set flanked with FseI (forward) and AgeI (reverse) RE sites. The PCR product was cloned in frame with an upstream His tag and downstream EGFP tag in a pCS2-FA vector; 6) pcDNA-E2F1; the ORF of human E2F1 was amplified using E2F1 cDNA [a gift from Doron Ginsberg (Bar Ilan University)] and a primer set flanked with KpnI (forward) and EcoRI (reverse) RE sites. The PCR product was cloned into pcDNA3.1(+) vector. The following E2F8 mutant variants were generated using pCS2-FA-E2F8 as a template: 1) <u>K</u>EN <u>m</u>utant E2F8-KM1 (^5^KEN^7^ to AAN); 2) KEN mutant E2F8-KM2 (^375^KEN^377^ to AAN); 3) KEN mutant E2F8-KM3 (^657^KEN^659^ to AAN); 4) RXXL mutant 1 (<u>D</u>-box <u>m</u>utant <u>1</u>) E2F8-DM1 (^87^RXXL^90^ to GXXV); 5) RXXL mutant 2 (D-box mutant 2) E2F8-DM2 (^183^RXXL^186^ to GXXV); 6) E2F8-DM1-DM2; 7) E2F8-KM1-KM2; 8) E2F8-KM1-DM2; 9) E2F8-KM2-DM2; 10) E2F8-KM3-DM2; 11) <u>D</u>NA-<u>b</u>inding <u>d</u>omain double mutant E2F8-DBD (^155^<u>RR</u>IYD^159^ and ^313^<u>RR</u>LYD^317^ to <u>AA</u>IYD and <u>AA</u>LYD); 12) <u>D</u>imerization <u>d</u>omain double mutant E2F8-DD (deletion of ^162^NVL^164^ and ^320^NVL^322^); 13) E2F8-DBD-DD (quadruple mutant); 14) E2F8-T20A (Thy 20 to Ala); 15) E2F8-T20D (Thy 20 to Asp); 16) E2F8-T44A (Thy 44 to Ala); 17) E2F8-T44D (Thy 44 to Asp); 18) E2F8-T20A-T44A (double mutant); 19) E2F8-T20D-T44D (double mutant); 20) <u>C</u>y <u>b</u>ox mutant <u>1</u> – E2F8-CB1 (^15^RGL^17^ to AGL); 21) <u>C</u>y <u>b</u>ox mutant <u>2</u> – E2F8-CB2 (^81^RGL^83^ to AGL); 22) <u>C</u>y <u>b</u>ox mutant <u>3</u> – E2F8-CB2 (^313^RRL^315^ to ARL); 23) <u>C</u>y <u>b</u>ox mutant <u>4</u> – E2F8-CB4 (^857^RKL^859^ to AKL). The following E2F8 variants were generated using pCS2-FA-His-E2F8-HA as a template: 1) E2F8-CB1; 2) E2F8-CB2; 3) E2F8-CB3; 4) E2F8-CB4. The following E2F8-N80 variants were generated using pCS2-His-E2F8-N80-EGFP as a template: 1) E2F8-N80-KM1 (^5^KEN^7^ to AAN); 2) E2F8-N80-T20A (Thy 20 to Ala); 3) E2F8-N80-T20D (Thy 20 to Asp); 4) E2F8-N80-T44A (Thy 44 to Ala); 5) E2F8-N80-T44D (Thy 44 to Asp); 6) E2F8-N80-T20A-T44A (double mutant); 7) E2F8-N80-T20D-T44D (double mutant). Overall, 40 new plasmids were generated. Aside from Tome-1, all ORFs were of human origin. All mutations were generated by site-directed mutagenesis (Agilent; #20521). Cloning and mutageneses were validated by Sanger sequencing. A full list of DNA oligos used for cloning is provided as supplementary information (**Table 1**).

### Cell culture

HeLa S3 (S3), HEK293T were originally from the ATCC. 293-T-REx™ cells were a gift from Vamsi Mootha (Harvard Medical School). All cell lines were cultured in Dulbecco’s modified Eagles medium (DMEM) supplemented with 10% fetal bovine serum, 2 mM L-Glutamine and 1% penicillin-streptomycin solution. Cells were maintained at 37°C in a humidified 5% CO_2_-containing atmosphere. S3 cells were either cultured on plates or in 1L glass spinner flasks in suspension (80 rpm). 293-T-REx™ cells were cultured in the presence of 5 µg/mL blasticidin (Gibco; #A11139-03) to maintain the pcDNA™6/TR plasmid carrying the ORF for tetracycline (Tet) repressor. The derivation of the Cyclin F knockout cell lines using CRISPR/Cas9 gene editing technology was previously described (Choudhury et al., 2016).

### Generation of <u>n</u>on-<u>d</u>egradable cyclin <u>B</u>1 cell line (NDB)

293-T-REx™ were stably transfected with pcDNA™4/TO plasmid carrying an open reading frame (ORF) of a <u>d</u>estruction-box <u>m</u>utant (DM) Cyclin B1 (Arg 42 and Leu 45 were substituted with Gly and Val, respectively) and a zeocin resistance gene. The ORF of cyclin B1-DM was amplified by PCR using the pCS2-FA-Cyclin B1-DM plasmid (Pe’er et al., 2013) as a template and a primer set flanked with RE sites for BamHI (forward) and XhoI (reverse). Zeocin concentration for selection: 200 µg/ml (BioBasic; # Z706211). NDB cells were stably transfected with a plasmid carrying the ORF of Histone H2B-YFP and a neomycin resistance gene. Stably transfected cells were selected by 500 µg/ml G418 (Formedium; #108321-42-2). NDB and NDB-H2B-YFP cell lines were originated from a single cell.

### Generation of inducible Cyclin F cell line

Cyclin F was cloned by gateway recombination cloning into pINDUCER20 (Meerbrey, Hu et al., 2011). Lentiviral particles were produced in HEK293T cells by transfecting cells with pINDUCER20-Cyclin F, as well as separate plasmids containing TAT, REV, VSVg and gag-pol. Viral particles were used to transduce MCF7 and T47D cells, which were subsequently selected with geneticin (Gibco #10131-035). Cyclin F expression was induced by the addition of doxycycline (5 ng/mL, 25 ng/mL, or 100 ng/mL) to the media for 48 hours and cells were analyzed by immunoblot. For *in vivo* experiments, MG132 (UBP Bio; #F1101) and bortezomib (Sigma-Aldrich; #5043140001) were used at a final concentration of 50 μM and 100 nM, respectively.

### Cell synchronization

#### Synchronization of S3 cells for mitotic extracts

A 400-500 ml culture of S3 cells was grown in suspension (1 L spinner flask, 85 rpm) until population reached a density of about 2.5 × 10^5^ cells/ml. Cells were then treated with 2 mM thymidine (Sigma-Aldrich; #T9250) for 22 h, washed and released into pre-warmed fresh media. After 3 h, cells were incubated with 50 ng/ml nocodazole (Sigma-Aldrich; #M1404) for 11-12 h, and harvested for mitotic extract preparation.

#### Synchronization of S3 cells for G1 extracts

Nocodazole-arrested cells (see previous paragraphed) were washed, recultured in 400-500 ml pre-warm fresh media, and harvested after 3.5 h for G1 extract preparation. Cells extracts were also generated from cells 6 and 8.5 h after release from nocodazole block.

#### Synchronization of S3 cells for S-phase extracts

S3 cells were cultured in suspension until population reached a density of approximately 5 × 10^5^ cells/ml. Cells were then incubated with 2 mM thymidine for 22 h, washed and released into pre-warmed fresh media for 9 h, and blocked again with 2 mM thymidine for 19 h. Cells were either harvested for extract preparation or released from the second thymidine block for 3 or 6 h and then harvested.

#### Synchronization of NDB cells in late mitosis

NDB cells were cultured in 15 cm/diameter plates. After reaching a confluency of about 75%, the cells were treated with 1 µg/ml tetracycline (Sigma-Aldrich; #87128) for 22 h and harvested for extract preparation or any other purpose.

#### Synchronization of NDB cells in pro-metaphase

NDB cells at 75% confluency were treated with 100 ng/ml nocodazole for 18 h.

#### Synchronization of NDB cells in G1

Asynchronous NDB cells showing the lowest 10% forward scatter width (FSC-W) signal were isolated by FACSAria III (BD). This sorting protocol yields a nearly pure G1 population without pre-synchronization (Vecsler, Lazar et al., 2013).

#### Synchronization of S3 cells for Western blotting

S3 cells arrested by thymidine-nocodazole block or double thymidine block (see above) were washed and either harvested (t = 0) or recultured for 30 min to 11 h, and then harvested.

### Live cell imaging

For light phase images, we used a Nikon eclipse TS100 inverted microscope equipped with x 20 (NA: 0.4) and x 40 (NA: 0.55) LWD lenses, a Nikon Digital-Slight DS-Fi1 camera, and a Nikon C-HGFI Intensilight. Images were processed by ImageJ software. Spread chromosomes were visualized using a Nikon Eclipse Ti-E inverted microscope equipped with x100 oil lens (NA: 1.4) or x40 Oil lens (NA: 1.3), Lumencor illuminator LED light source, and a Zyla sCMOS camera (Andor Technology). Filter sets: excitation 390/18 nm; emission 460/50 nm. Images were processed by Nikon NIS-element and ImageJ software. Time-lapse microscopy of NDB was performed with a Leica SP8 inverted scanning confocal microscope equipped with an x63 oil lens (NA: 1.4), and HyD detector. Excitation: 488 nm laser. Emission: 511-552 nm. Images were acquired and processed using *LASX and ImageJ software*.

### Flow cytometry

Overall, cell synchronization was measured by DNA quantification following a standard propidium iodide (PI) staining protocol (Sigma-Aldrich; #81845). Gallios (Beckman Coulter) and FACSAria III (BD) flow cytometers were used for analyzing the stained cells. Cell cycle phase distribution was determined by ModFit LT™’s Sync Wizard model (Verity Software House). FACSAria III was also used to sort G1 NDB cells for Western blot analysis (see above) and for sorting single NDB and NDB-H2B-YFP cells.

### Chromosome spread

Tet-induced- and nocodazole-arrested NDB cells (see above) were harvested by gentle pipetting, washed gently with PBS, lysed in hypotonic solution (0.8% KCl, 10 min at room temperature) and fixed in a methanol/glacial acetic acid solution (3:1 volume ratio). Cell droplets were released from one-meter height onto tilted glass slides. The slides were air-dried and mounted with a mounting solution (Thermo Fisher Scientific; 4112APG) and DAPI stain [5 μg/ml (Sigma-Aldrich; #P9542)].

### Western blotting

#### Protein lysis

Cells were washed twice in cold PBS and lysed in a cold lysis buffer (50 mM Tris pH 7.6, 150 mM NaCl, 5 mM EDTA pH8.0, 0.5% NP-40) supplemented with a protease inhibitor cocktail (Roche; #4693159001), 1 mM <u>p</u>henyl<u>m</u>ethyl<u>s</u>ulfonyl fluoride (PMSF), phosphatase inhibitor cocktail (Sigma-Aldrich; #P5726, #P0044), 10 mM NaF; 20 mM β-glycerophosphate; 1 mM Na_3_VO_4_, 20 mM <u>P</u>-<u>n</u>itro<u>p</u>henyl<u>p</u>hosphate (PNPP). Protein extracts were incubated on ice for 30 min and the non-soluble components were pelleted by centrifugation for 45 minutes at 14,000 x g. Protein concentration was determined by a standard Bradford assay (Bio-Rad #500-0006), a linear BSA calibration curve, and an Epoch microplate spectrophotometer.

#### Immunoblotting

Protein samples were mixed with Laemmli buffer, denatured (5-10 min, 95°C) and resolved on freshly-made 8-10% acrylamide gels using a Tris-glycine running buffer. Proteins were then electro-transferred onto a nitrocellulose membrane (Bio-Rad; #162-0115) using a wet transfer or Trans-Blot Turbo™ transfer system (Bio-Rad). Transfer quality was verified by Ponceau S Solution (Sigma-Aldrich; #81462). Membranes were washed, blocked (5% skimmed milk in TBST) and incubated overnight (4°) with antibody solution (2.5% BSA and 0.05% Sodium Azide in PBS). The following primary antibodies were used: anti-Tubulin (DSHB; #12G10,), anti-Actin (DSHB; #JLA20), anti-HSP70 (Santa Cruz Biotechnology; #SC-24), anti-GAPDH (Santa Cruz Biotechnology; #SC-47724), anti-E2F8 (Abnova; #H00079733-M01 and Abcam; #AB109565), anti-E2F1 (Santa Cruz Biotechnology; #SC-193), anti-Securin (Abcam; #AB3305), anti-Geminin (Abcam; #AB12147), anti-Kifc1 (Bethyl Laboratories; #A300-951A), anti-Cyclin B1 (Santa Cruz Biotechnology; #SC-70898) anti-Cdc20 (Santa Cruz Biotechnology; #SC-8358), anti-Cdh1 (Calibochem; #CC43-100UG), anti-Cdc27 (Santa Cruz Biotechnology #SC-5618 and SC-9972), anti-Cyclin F (Santa Cruz Biotechnology; #SC-952), anti-HA (Biolegend; #901502); anti-FLAG (Sigma; #A8592). Horseradish peroxidase-conjugated (HRP) secondary antibodies were purchased from Jackson ImmunoResearch: #115-035-174; #115-035-144; #115-035-003). ECL signal was detected by a SuperSignal West Femtochemiluminescence substrate (Pierce; #34095) or an EZ-ECL (Biological Industries; #20-500-171).

### Immunoprecipitation

APC/C was immunoprecipitated from NDB mitotic extracts pre- and post 30 min incubation with RO-3306 (see above). To this end, 200 μg cellular extracts were mixed with 300 μl wash buffer (150 mM NaCl, 20 mM Tris HCI pH=7.5, 10% glycerol, 0.1% Triton, EDTA 1 mM) and protease inhibitor cocktail (Sigma-Aldrich; #p2714-1BTL). Next, 15 μl agarose-conjugated anti-Cdc27 antibodies (Santa Cruz Biotechnology; SC-9972AC) were incubated with the diluted extracts for 4 h at 4°C. Beads were washed twice with 150 mM NaCl wash buffer and once with 75 mM wash buffer, resuspended with 4x Laemmli sample buffer and denatured 10 min at 96°C. Protein samples were resolved by SDS-PAGE.

#### HA/FLAG immunoprecipitation

HEK293T cells were transfected with the indicated plasmids using PolyJet DNA *in vitro* transfection reagent (SignaGen #SL1000688). 24 hours post-transfection, cells were harvested, lysed in NETN buffer [20 mM Tris, pH 8.0, 100 mMN NaCl, 0.5 mM EDTA, 0.5% NP40, 2 µg/mL pepstatin, 2 µg/mL apoprotinin, 10 µg/mL leupeptin, 1.0 mM 4-(2 aminoethyl) benzenesulfonyl fluoride, 1.0 mM Na3VO4] and immunoprecipitated with EZview Red anti-HA or EZview Red anti-FLAG M2 affinity gel (Sigma #E6779; #F2426). IPs were performed for 2 hours at four degrees, washed 5 times in cold NETN buffer by rotating for 5 minutes each time, and eluted in Laemmli buffer, prior to analysis by immunoblot.

### Preparation of cell extracts

#### HeLa S3 extracts

Synchronous S3 cells were washed with cold PBS and lysed in a swelling buffer [20 mM Hepes pH 7.5, 2 mM MgCl2, 5 mM KCl, 1 mM DTT, and protease inhibitor cocktail (Roche; #11836170001)] supplemented with energy regenerating mixture (1 mM ATP, 0.1 mM EGTA, 1 mM MgCl2, 7.5 mM creatine phosphate, 50 μg/ml creatine phosphokinase). Cells were swelled on ice for 30 min and homogenized by freeze-thawing in liquid nitrogen and passed through a 21G needle 10 times. Extracts were cleared by subsequent centrifugations (14,000 RPM; 10 and 40 min), and stored at −80°.

#### NDB mitotic extracts

Tet-induced NDB cells were harvested from twenty 150 mm plates. Cells were washed gently with cold PBS and lysed for extract preparation (see previous paragraph).

#### NDB G1-like extracts

NDB mitotic extracts (see above) were pre-activated with RO-3306 for 15 to 30 min. This treatment overrides the blocking effect of non-degradable Cyclin B1, and effectively induces mitotic exit into an APC/C^Cdh1^-active state in tube. Only then *<u>i</u>n <u>v</u>itro* <u>t</u>ranslated (IVT) substrates were added to the reaction mixture for degradation assays.

### Degradation and mobility shift assays

Target proteins were *in vitro* transcribed and translated in rabbit reticulocyte lysate (TNT-coupled reticulocyte system; Promega; #L4600, #L4610) supplemented with ^35^S-Methionine (IsoLabel L-35S Steady Blue, Izotop; #TSM-01). Degradation assays were performed in 20 μl cell extracts supplemented with 1 μl of x20 energy regenerating mixture (see above), 1 μl of 10 mg/ml Ubiquitin (Ub) solution (Boston Biochem; #U-100H) and 1 µl radiolabeled IVT product of interest. As indicated, reaction mixture were supplemented with one or more of the following reagents: 1) recombinant His-tagged UbcH10 or UbcH10^DN^ (5 μg); 2) recombinant GST-tagged Emi1 C-terminus (1 μg); 3) Cdk1 inhibitor RO-3306 [(15-30 μM); Enzo Life Sciences; #ALX-270-463-M001]; 4) MG132 [(20 μM); Boston Biochem; #I-130)]; and 5) DMSO. Reaction mixture were incubated at 28°C, unless otherwise is indicated. Aliquots of 3-5 μl were taken every 10 to 20 min, mixed with Laemmli buffer, denatured (10 min, 95°C), and were quickly frozen in liquid nitrogen. Protein samples were resolved by SDS-PAGE. Gels were soaked in a methanol/acetic acid (10%/7.5%) solution for 20 min, dried in vacuum and heat, and exposed to phosphor screen. IVT proteins were visualized by autoradiography using Typhoon™ FLA 9500 phosphorimager (GE Healthcare Life Sciences). Signal intensity was measured by ImageJ software after background subtraction, and normalized to ^35^S signal at time 0 min (t_0_). All plots were created in Microsoft Excel software, version 16.20. Mean and standard error (SE) were calculated from three or four independent degradation assays.

### On-chip ubiquitination and Ub-chain preference assays

#### Mold Fabrication

The device was designed using AutoCAD 2011 (Autodesk, Inc.) and each layer was reproduced as a chrome mask at 40,000 dpi (Fineline-Imaging). Flow molds were fabricated on 4” silicon wafers (Silicon Quest International) with pretreatment of O_2_ plasma 34% for 5 min. The wafers were spin-coated with SPR 220-7 (Shipley; 1500 rpm, 60 sec) yielding a substrate height of around 13-15 μm. The molds were baked at 105°C for 6 min followed by a 120 sec I-line exposure on a MA6 contact mask aligner (Karl Suss). Next, molds were incubated for 2 hrs in RT, baked in 110°C for 5 min, incubate for additional 45 min at RT, and developed with AZ 726 MIF Developer followed by DW H_2_O wash. Finally, molds were annealed at ramping temp (70-200°C; 10°C/h) for 15 hrs. Control molds were fabricated on 4” silicon wafers by spin coating SU-8 2025/3025 (MicroChem) at 500 rpm for 5 sec followed by 3000 rpm for 70 sec yielding a substrate height of around 15-18 μm. The molds were baked at 65°C for 2 min and 95°C for 7 min. Next, the wafers were exposed for 15 sec on the mask aligner, followed by a post-exposure baking series of 65°C for 1 min and 95°C for 3 min. The wafers were developed in AZ EBR Solvent for 4.5 min followed by an isopropanol wash. At the end of the fabrication step, control and flow molds were Teflon coated to promote elastomer release during following use.

#### Device Fabrication

The microfluidic devices were fabricated on silicone molds casting silicone elastomer polydimethylsiloxane (PDMS, SYLGARD 184®, Dow Corning). Each device consists of 2 aligned PDMS layers: the flow and the control layer. A mixture of a silicone-based elastomer and curing agent was prepared in 2 different ratios 5:1 and 20:1 for the control and flow molds, respectively. The control layer was degassed and baked for 30 min at 80°C /60 min at 90°C. The flow layer was initially spin coated (Laurell Technologies) at 1500-2000 rpm for 60 sec and then baked at 80°C for 30 min. The control layer was separated from its mold and then control channel access holes were punched. The flow and control layers were aligned manually and baked for 2 h at 80°C. The 2-layer device (chip) was peeled from the flow mold and flow channels access holes were punched.

#### Surface Chemistry

In order to bind the expressed protein, we first cover the epoxy slides with biotinylated BSA (1 µg/µl, Thermo Scientific). Streptavidin (0.5 µg/µl Neutravidin, Pierce) was then introduced, to interact with the biotinylated BSA layer. Next, a designated set of pneomatic valves, also known as ‘button valves’ (Noach-Hirsh et al., 2015), were closed and a layer of biotinylated PEG (1 µg/µl, Nanocs) was used to block the protein chmaber periphery, leaving the center itself exposed to avidin to which antibodies and eventually target proteins can be attrached. Then, the button valves were opened and anti-GFP biotinylated antibodies (0.2 µg/µl, Abcam; #ab6658) were flowed through the device to interact with the exposed avidin at the protein chamber center. This creates an array of protein chamber with anti GFP antibodies to which EGFP-tagged E2F8 IVT product can bind. PBS buffer or PBS buffer with 1% BSA was used to the wash between steps.

#### On-chip ubiquitination and ubiquitin-chain preference assays

E2F8-EGFP IVT product was flowed into the chip and immobilized on the surface at the protein chamber via its EGFP tag, followed by a wash with a PBS buffer. S3 G1 extracts were mixed with i) 0.04 mg/ml <u>Rh</u>odamin-coupled <u>Ub</u>iquitin (Rd-Ub) (Boston Biochem; #U-600); ii) mock or 0.4 mg/ml of one of the following <u>unlabled</u> Ubiqutin variants: WT (Boston Biochem; #U100-H), Lys11 to Arg mutant (Boston Biochem; #UM-K11R), Lys48 to Arg mutant (Boston Biochem;#UM-K48R), Lys63 to Arg mutant (Boston Biochem; #UM-K63R); and iii) mock or 0.3 mg/ml His-tagged UbcH10^DN^ (recombinant) or 0.48 mg/ml GST-tagged Emi1 C-terminus (recombinant). Next, reaction mixture were flowed to protein chambers for 10 min (RT). Unbound material was washed by an Hepes buffer (50 mM). Rd-Ub levels were determined by 535 nm excitation (emission filters: 575/50). Rd-Ub signal was normalized to the immobilized protein level, as measured by 488 nm-excited GFP (emission filter: 535/25). This normalization provides net ubiquitnation signal per micro-compartment. Ubiquitin-chain preference of E2F8 was validated in tube using degradation assays in S3 G1 extarcts to which 8 µg UbcH10 (recombinant), and 0.4 µg WT- or mutant K-to-R Ub (see above) were added.

#### Image and Data Analysis

LS Reloaded microarray scanner (LS Reloaded, Tecan, Männedorf, Switzarland) and GenePix7.0 (Molecular Devices) image analysis software were used for analysis and data esentation.

For more details, see Noach-Hirsh et al. (Noach-Hirsh et al., 2015).

## Supplementary Information

**Figure S1:**
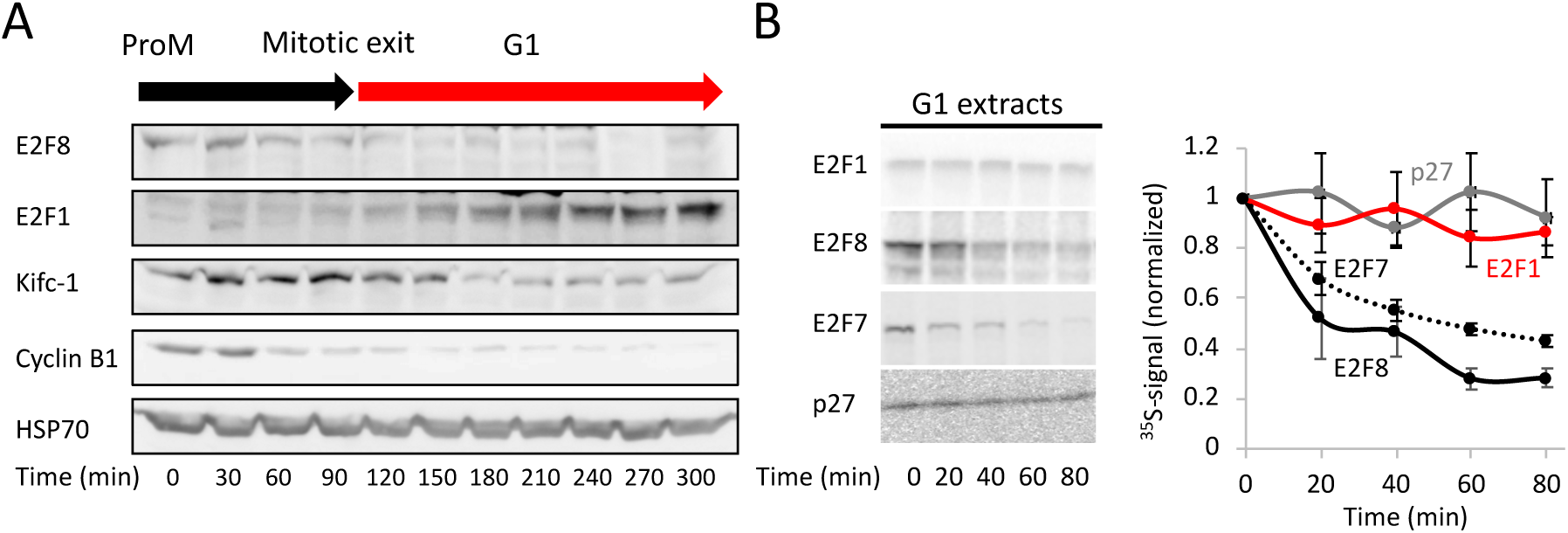
E2F1 is stable in APC/C^Cdh1^-active G1 extracts. **(A)** Western blot analyses of E2F1, E2F8, Cyclin B1, Kifc-1 (APC/C^Cdh1^ targets) and HSP70 (loading control) in synchronous S3 cells released from a thymidine-nocodazole block into G1. Samples were harvested in 30 min intervals. **(B)** Time-dependent degradations of ^35^S-labeled E2F1, E2F7, E2F8 and p27 (IVT products) in G1 extracts were assayed by SDS-PAGE and autoradiography. A set of source data and quantification of three experiments are shown (mean and SE values are plotted). In contrast to E2F8 and E2F7, E2F1 is stable in G1 extracts, like the SCF^Skp2^ target p27 (negative control).

### Degradation of E2F8 in G1 extracts is dependent on intact dimerization domains

E2F8-EGFP, but not EGFP-E2F8, is degraded in G1 extracts in a manner similar to untagged E2F8 (**Fig. S2**). At some point during this study, we asked to complement our findings with *in vivo* quantification of fluorescently tagged E2F8 in live cells. Multiple attempts and strategies to generate stable cell lines with a constitutive (not inducible) expression of E2F8-EGFP failed. In addition, extensive efforts to knock in Venus (YFP) into an endogenous E2F8 locus also failed. Reasoning that fluorescently tagged E2F8, all the more so when overexpressed, might be cytotoxic, and focusing on E2F8 dynamics rather than function, we generated E2F8 lacking <u>D</u>NA <u>b</u>inding <u>d</u>omains (DBD1/2) with the assumption that dysfunctional E2F8-EGFP might be inert *in vivo*. Because E2F8 functions as a homo/hetero-dimer, we also deleted its two <u>d</u>imerization <u>d</u>omains (DD1/2) to minimize potential dominant-negative effect of the modified E2F8 in cells (**Fig. 5A and Fig. S3A**). Mutations were strategized based on the literature (Liu, Shats et al., 2013, Zalmas, Zhao et al., 2008). Conceptually, this experimental set up is valid only if the temporal proteolysis of the modified E2F8 variants is unchanged. To test that, we first assayed the proteolysis of the quadruple DBD/DD E2F8 mutant in G1 extracts. Surprisingly, this mutation nearly blocked E2F8 degradation (**Fig. S3B**). It was the DD mutations, rather than the DBD mutations, that contributed most to this molecular phenotype. In view of these results, we decided to abandon this line of research. The data, however, have implications on the structure-to-function relationship of E2F8 with relevance to past and future research of atypical E2Fs.

**Figure S2:**
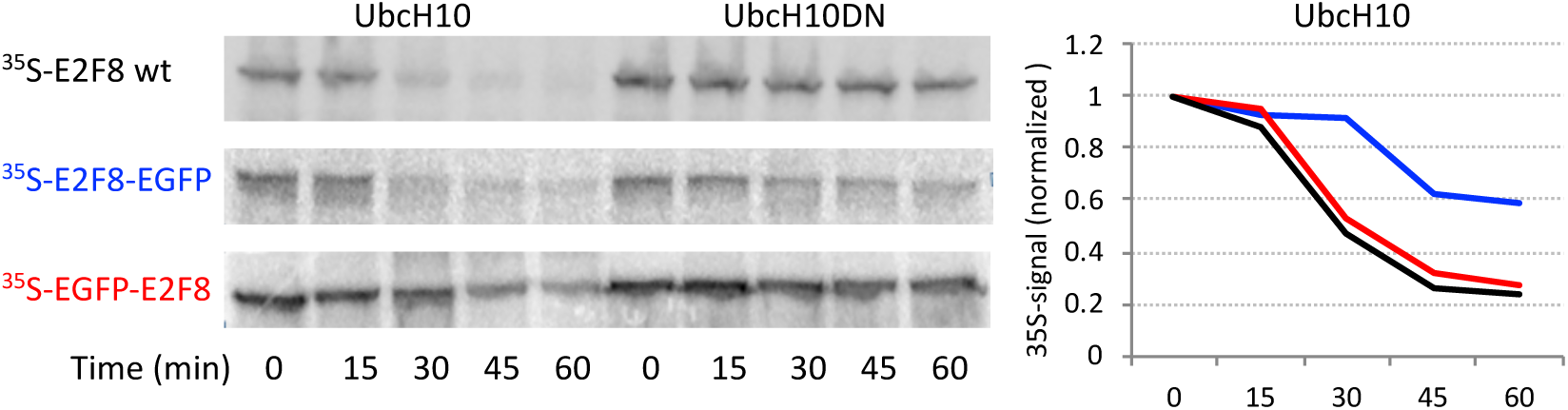
Proteolysis of EGFP-tagged E2F8 in G1 extracts. E2F8 was tagged with EGFP on either N- or C terminus. ^35^S-labeled IVT product of tagged and untagged E2F8 were made, and their time-dependent degradations in G1 extracts were assayed by SDS-PAGE and autoradiography. Quantification of source data is plotted using matching colors.

**Figure S3.**
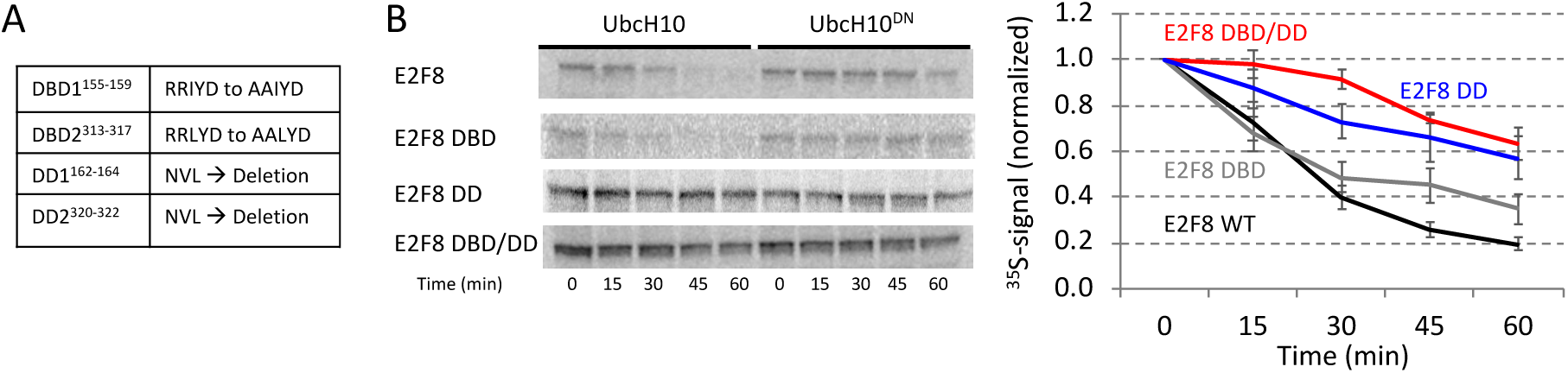
Degradation of E2F8 in G1 extracts is dependent on intact dimerization domains. **(A)** E2F8 variants with mutated <u>D</u>NA-<u>b</u>inding <u>d</u>omains (DBD1/DBD2) or dimerization domains (DD1/DD2) or both, were generated using point mutations or deletion. **(B)** Degradation of ^35^S-labeled E2F8 variants (IVT products) was tested in G1 extracts supplemented with WT or dominant negative UbcH10 (UbcH10^DN^). Time-dependent degradation was assayed by SDS-PAGE and autoradiography. Representative raw data and quantifications are shown. Mean E2F8 levels (^35^S signals) normalized to max signal at *t* = 0 are shown (*n* = 3-4). Bars represent SE. An unexpected, albeit profound, dependency of E2F8 proteolysis on intact dimerization domains, but not the adjacent DNA-binding domains, is evident.

### Supplementary Tables

**Table S1:**
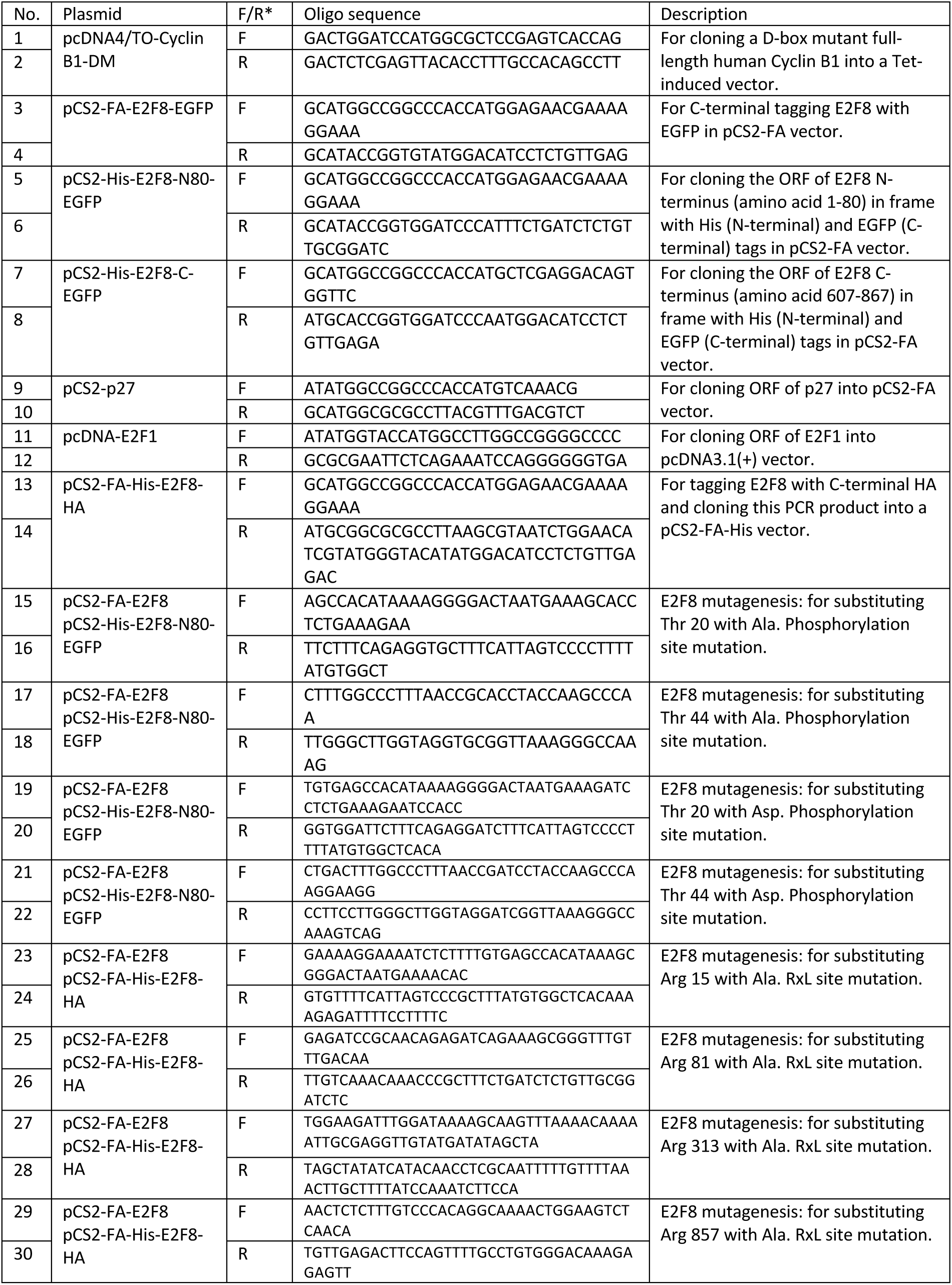
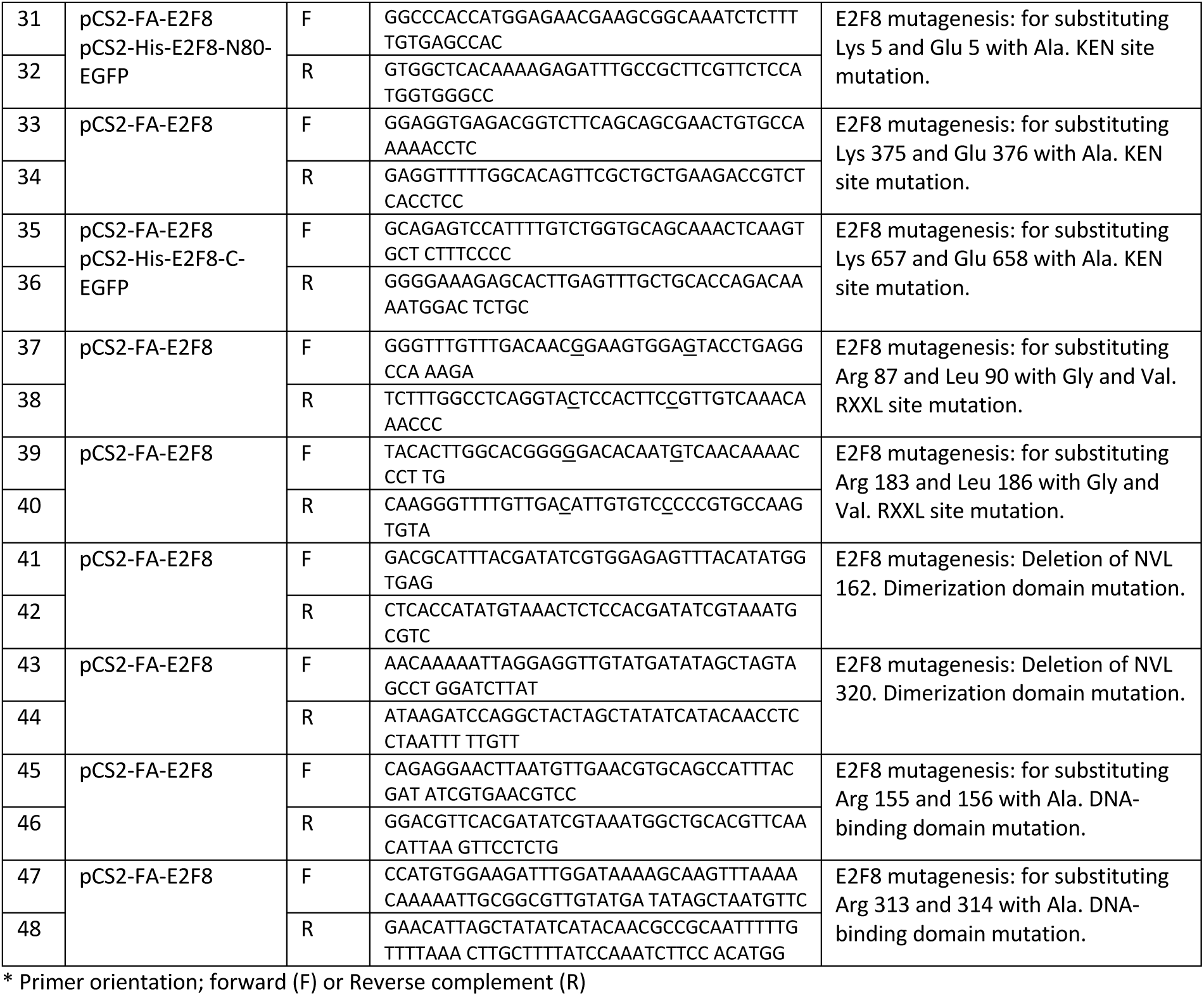
List of DNA oligos used for cloning and mutagenesis

